# Co-expression Patterns Explain how a Basic Transcriptional Role for MYC Modulates *Wnt* and MAPK Pathways in Colon and Lung Adenocarcinomas

**DOI:** 10.1101/2021.10.28.466287

**Authors:** Melanie Haas Kucherlapati

## Abstract

Genome duplication begins at many epigenetically determined sites by pre-replication, pre-initiation, and replisome complexes; co-expression of their components must be optimally timed for S phase to occur. Oscillations of cyclin dependent kinases (Cdks) and regulator cyclins control cell cycling, many are pharmacological targets in cancer. This study examines gene expression relationships between drivers, cell cycle components, and a subset of proliferation genes in colon (COAD) and lung (LUAD) adenocarcinomas. Several known drivers of COAD and LUAD including APC, CTNNB1, KRAS, MYC, Braf, TP53, Rb1, and EGFR are also observed with focus on *Wnt* and MAPK signaling activation. *Wnt* signaling activation has relevance for immune checkpoint inhibitor therapy, as it provides cancer cells with escape mechanisms.

MYC and KRAS co-expressed directly with far fewer proliferation genes in LUAD than COAD, suggesting their expression is ectopic to S phase in lung tumors. APC indirectly co-expressed with the same factors in both COAD and LUAD, but was found co-expressed indirectly with MYC and mutated only in COAD. Other *Wnt* signaling components also co-expressed in low MYC context in COAD, had significantly higher mutation frequencies. These data suggest *Wnt* signaling activation to be the indirect result of decreased MYC availability in COAD, and ectopic overexpression of MYC in LUAD. Cyclins CCNH, CCNC, and CCNK, co-expressed with far fewer proliferation genes in LUAD. Conversely, Braf had direct co-expression with many proliferation factors in non EGFR activated LUAD. Proliferation in EGFR activated LUAD was completely deregulated with E2F(s) 4/5/6 expression, potentially explaining their low proliferative ability.

## Background

The adenomatous polyposis coli (APC) tumor suppressor gene is mutated in 70-80% of all COAD cases [1]. APC controls proliferation through MYC which is normally expressed in S phase [2]. The resulting dysregulation of the *Wnt* signaling pathway is facilitated by changes to the cytoskeleton regulated mostly by APC truncation mutation and CTNNB1 activation. A body of evidence also implicates the *Wnt* signaling pathway in LUAD development and progression. Model systems have shown that the pathway is essential for initiation and maintenance of Braf driven tumors [3] [4]; KRAS and Braf tumor progression occurs with *Wnt* pathway activation [5]. Metastasis has been associated with increased *Wnt* expression [6] [7]. However APC mutation happens with far less frequency in LUAD than it does in COAD [8, 9].

Although proliferation of both COAD and LUAD involves deregulation of the *Wnt* signaling pathway, the canonical pathways taken by the two tumor types seem to be different. To better understand how the pathways are modulated in these two tumor types Spearmen’s co-expression correlations (see Materials and Methods) have been used to examine the relationship of proliferation gene expression with the expression of selected cell cycle components and drivers involved in *Wnt* and MAPK signaling pathways. Identification of *Wnt* signaling modulation has relevance for immune checkpoint inhibitor therapy, as its aberrant activation provides escape mechanisms.

As expected MYC expression has a clear direct relationship to proliferation gene expression in COAD, and an indirect relationship with APC. Unexpectedly, while APC had an indirect relationship to proliferation gene expression in LUAD, MYC/MYCL/MYCN and KRAS directly co-expressed with far fewer replication genes and did not have a significant indirect expression relationship with APC. Universal Cdk inhibitor CDKN1A (p21 Cip/Waf1) whose expression is normally turned off by MYC and whose protein competes with MYC for a PCNA binding site, also did not indirectly co-express with MYC in LUAD as it does in COAD supporting deregulation of MYC expression in LUAD. CDKN1A does significantly indirectly rank with Braf in LUAD, and Braf direct co-expression with proliferation factors was found elevated in non-EGFR activated LUAD. In normal lung and colon MYC/MYCL/MYCN and KRAS expression are comparable. That MYC co-expression with proliferation factors does not rank directly with high frequency suggests that MYC expression is deregulated with respect to S phase. In addition to regulating many genes, MYC is also thought to function in basic transcriptional elongation, a role which could explain how APC comes to be mutated in COAD but not in LUAD and have ramifications, with APC as a paradigm, for why it has been so difficult to find drivers in some LUAD.

MYC is a transcriptional transducer of *Wnt* signaling, as is Mediator complex [10]. Mediator of RNA polymerase II transcription complex includes, among other proteins, CCNC: Cdk8/19, and Med12/13. CCNC co-expression ranked directly with a high frequency of proliferation genes in COAD, less in LUAD; unexpectedly, Cdk8 was not highly correlated with proliferation in COAD. Since Cdk8 overexpression occurs in about half of all COAD, it is thought of as an oncogene [11]. The present study indicates Cdk8 over-expression is not during S phase alone in COAD, and that not all cases with high Cdk8 expression necessarily have high proliferative potential. Components of CCNY: Cdk14, a membrane complex also conceptually linked to *Wnt* signaling but whose peak levels of expression occur during M phase [12], are found accordingly to anti-correlate with proliferation in both COAD and LUAD.

LUAD have been separated in this study into “non-EGFR” and “EGFR activated” groups; other known pathways of tumorigenesis are not observed. EGFR activated tumors (all non-treated) have relatively low proliferative capacity and appear to have stopped DNA synthesis as part of a DNA damage checkpoint [13], or as part of an embryonic diapause-like mechanism. In “EGFR activated” cases co-expression of the proliferative subset with E2Fs 1-8 shows canonical repressor E2Fs 4/5/6 co-expression are completely unsynchronized.

## Methods

### Tumor cohorts

The colon adenocarcinoma (COAD) tumor cohort used throughout this study was assembled and analyzed by The Cancer Genome Atlas network (TCGA) [1]. COAD clusters were defined as having “Chromosomal Instability” (CIN), “Genome stabile” (GS), “Microsatellite unstable” (MMR), or with “POLE “mutation. Lung adenocarcinomas (LUAD), also assembled and analyzed by TCGA [8, 9], were placed into six clusters based on copy number, DNA methylation, and mRNA expression. Clusters for both COAD and LUAD were identified in heatmaps for individual genes and their exons. All mutation, expression, and co-expression data generated by TCGA is available in cBioPortal [14] [15, 16], the Genome Data Commons [17] and the Broad Institute [18].

### Proliferation Genes Examined

This study examines thirty-one proliferation genes known to be associated with origin licensing, firing, and DNA synthesis (Table 1). The gene subset is used as a marker of replication potential; selection was based on function and included a substantial number of genes viewed as essential to non-neoplastic replication initiation and processing. Test genes are compared to each gene of the subset and the frequency of significant comparisons reported, 30% or above is arbitrarily highlighted.

**Table 1.**
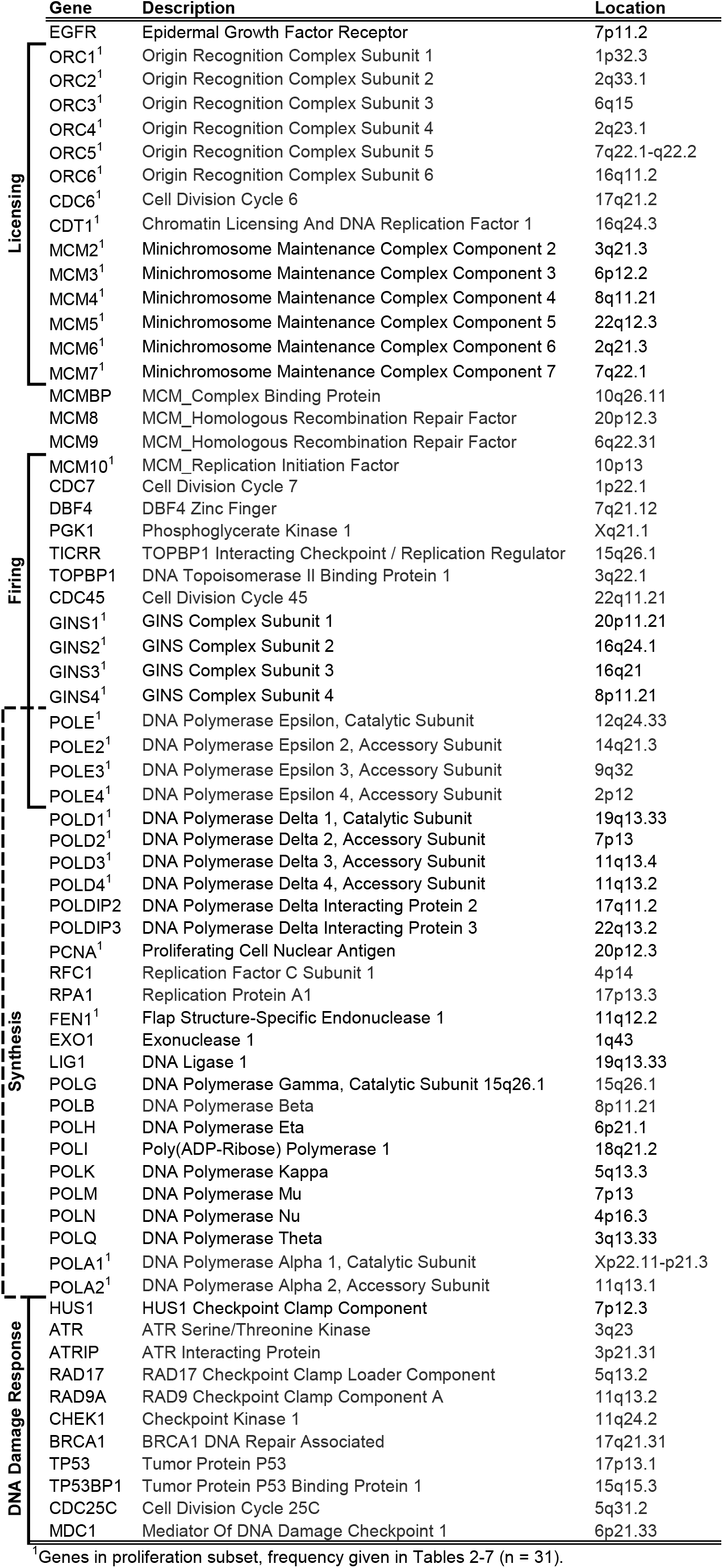
Selected Genes Associated with DNA Replication.

### Co-expression of Proliferation Genes with Cell Cycle Genes and Putative Drivers

Spearman’s correlation (r_s_) [19] is used to identify significant co-expression (mRNA, RNA Seq V2 RSEM) relationships identified as “direct” “indirect” or “none” found by the formula:

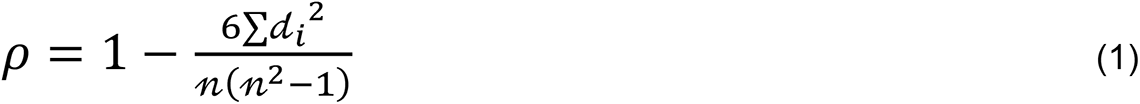

where *d_i_* = difference in paired ranks based on the expression data, and *n* = number of cases.

r_s_ (*ρ*) was calculated by cBioPortal [14], significance determined by two-sided t test with Benjamini Hochberg false discovery rate (FDR) correction procedure applied. Spearman’s correlation coefficient can take values from +1 to −1, where +1 indicates perfect association, zero indicates no association, and −1 indicates a perfect negative association. Lack of co-expression (strength and direction) upon ranking is taken to indicate that the levels of gene product in question are not coordinated with origin licensing, firing, or “normal” DNA synthesis. This could occur under two conditions, either the genes never were transcriptionally coordinated or they lost regulation of expression due to tumorigenesis. Both the Genotype-Tissue Expression (GTEx) Database [20] and relevant literature are used to discriminate one possibility from the other.

By way of clarification, many processes (e.g. DNA damage response, repair mechanisms, mitosis) closely associated with proliferation are not being measured by the subset. Only the co-expression relationship of a gene with the proliferation subset representing non-neoplastic S phase is presented, observations suggested by co-expression were validated with other cBioPortal tools (e.g. expression heatmaps) when necessary.

### Other Statistical Tests

Fisher’s exact test was performed [21] to determine significance of proliferation factors with direct versus no direction association between COAD and LUAD, for cyclin regulator genes CCNH, CCNK, and CCNC and MYC.

An unpaired t-test/two-tailed was performed [22] to determine significance in the means of C to T mutations, and APC/EGFR expression levels, between COAD and LUAD.

Chi square analysis with Yates correction was performed [21] (as values too large for Fisher’s exact), to determine significance of the number of mutations versus wildtypes in *Wnt* signaling genes (SMAD(s) 2-4, TCF7L2, APC, TP53, CSNK2A1, FZD1, CCND1) in COAD versus LUAD PANCAN cases.

## Results

Because of the complexity of cell cycling, in each segment of “Results” a brief statement is given explaining the relevant gene’s normal function prior to describing the study’s findings.

### Proliferation subset

A list of proliferation genes examined is shown in Table 1 (n = 66); thirty-one genes are selected from that list as a proliferation marker subset with most checkpoint and repair proteins intentionally excluded. The abbreviated list of thirty-one genes is subsequently referred to as the “subset”.

### Cyclin E: Cdk2/1, Cyclin D: Cdk4/6, and Rb1

Canonical cyclins have periodic expression during the cell cycle, they complex with specific Cdks controlling their kinase activity. During late G1 and S phase, CCNE1 and Cdk2/1 are highly expressed, CCND1:Cdk4/6 is reduced. In normal cells when CCND1:Cdk4/6 is expressed, activated, and appropriately distributed subcellularly, it is sensitive to extra-cellular growth factors and controls progression through G1, and the G1 to S phase transition [23].

In this study all components of the CCNE1:Cdk2/1 complex directly co-expressed with 94-97% of the genes in the subset in both COAD and LUAD (Tables 2-5; Supplementary Figure 1). Overall alteration in expression and mutation to CCNE1 was 7% in COAD, and 5% in LUAD [14–16].

**Table 2.**
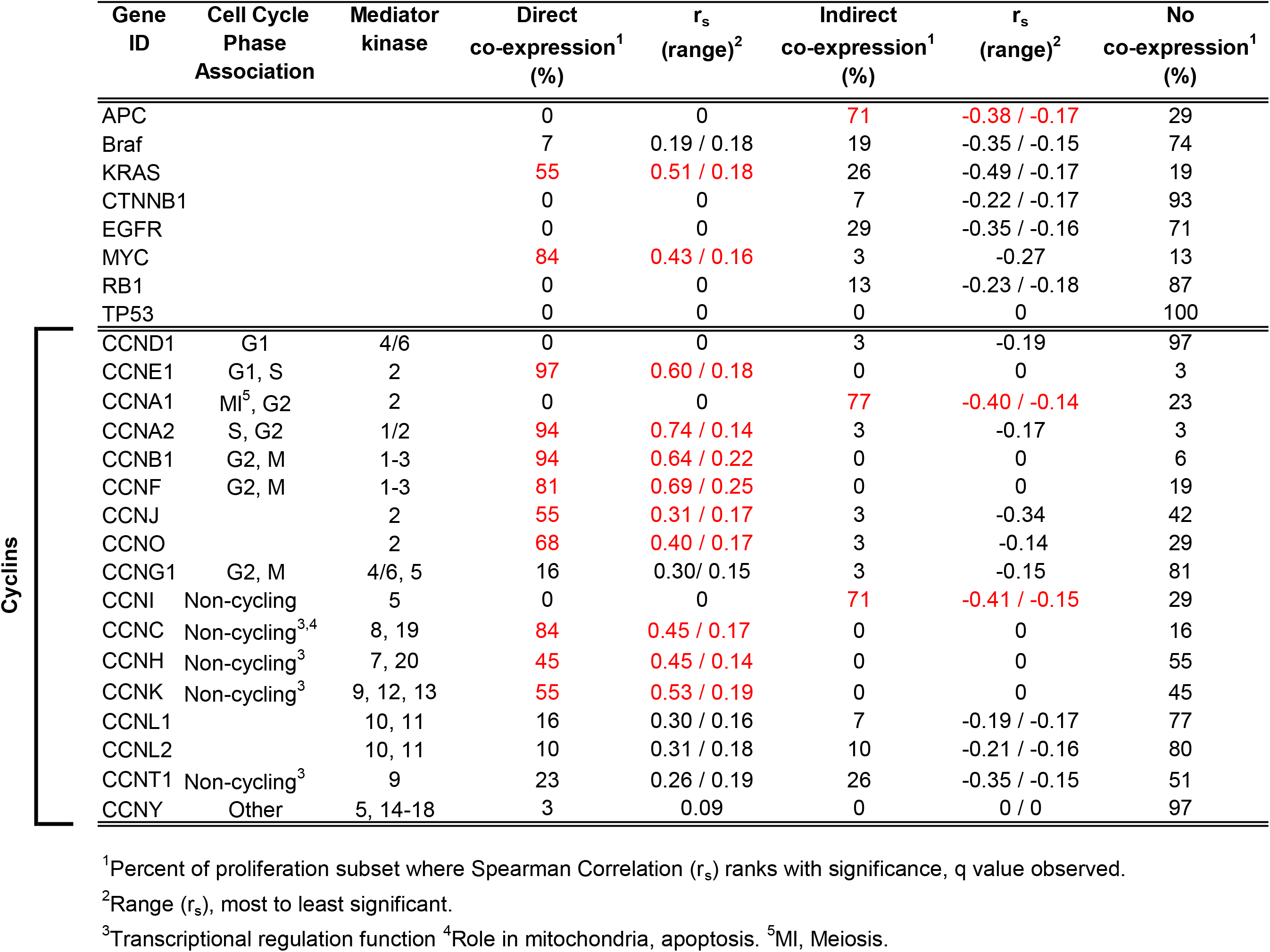
Co-expression of Proliferation Genes with Drivers and Cyclins in COAD (n = 276)

CCND1 had no association with most proliferation factors in COAD or LUAD; overall alteration was 4% in both tumor types. While Cdk4 co-expression ranked directly with a high frequency of proliferative components in both tumor types, CCND1 and Cdk6 did not (Tables 2-5).

When LUAD were examined as “non-EGFR” versus “EGFR protein kinase activated” cases, CCNE1 directly co-expressed with 97% of the subset in non-EGFR tumors only (Table 6).

CCND1: Cdk4/6 proteins physically interact with the retinoblastoma protein, the resulting phosphorylated complex regulates transcription patterns through E2F factors [24, 25]. Rb1 did not have significant co-expression with the subset in COAD (Table 2), nor was it mutated or aberrantly expressed. Rb1 indirectly co-expressed with a low frequency of the subset in LUAD (Table 3); overall alteration was 13%.

**Table 3.**
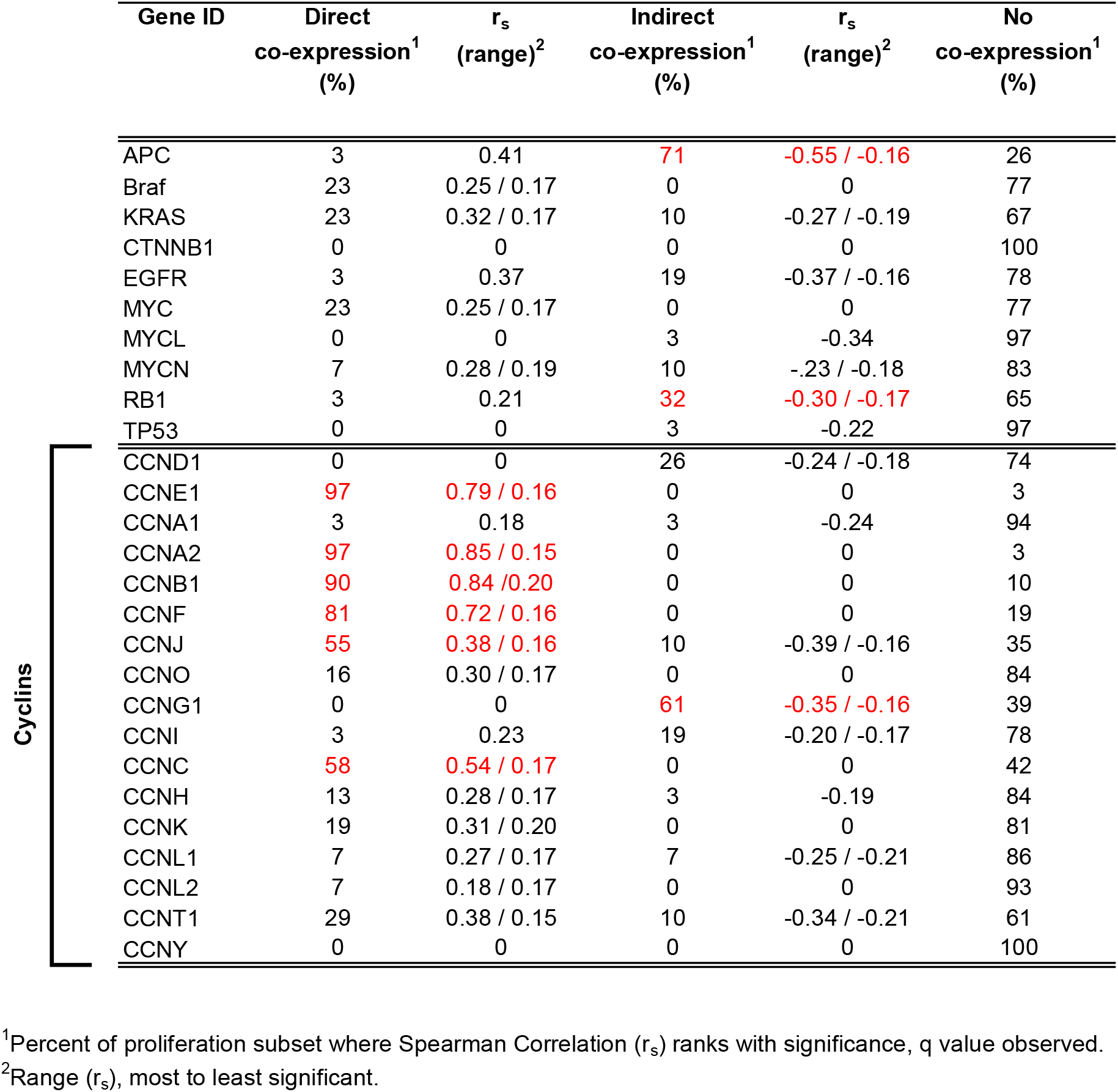
Co-expression of Proliferation Genes with Drivers and Cyclins in LUAD (n=230)

### CCNA2: Cdk1/2, CCNA1:Cdk2, CCNB1:Cdk1, and TP53

CCNA2: Cdk1/2 functions in both G1/S transition and down-regulation of G2/M through E2F factor phosphorylation [26–28]. Components of CCNA2:Cdk1/2 co-expressed directly with 94-97% of the subset in both tumor types (Tables 2-5).

CCNA1: Cdk2 is associated with the meiotic cell cycle [29–31]; the cyclin co-expressed indirectly with some of the subset in COAD, not LUAD (Tables 2-3).

Multi-functional tumor suppressor TP53 was mutated and/or aberrantly expressed in both COAD (56%) and LUAD (48%), there was no co-expression association with the proliferation subset in either tumor type (Tables 2-3).

CCNB1: Cdk1 (maturation-promoting factor, MPF), is essential for G2/M transition [32]. Both regulator and kinase directly co-expressed with over 90% of the proliferation gene subset in both tumor types (Tables 2-5).

### Atypical CCNF

Protein degradation provides an additional level of regulation to the cell cycle; CCNF is part of the Skp1-Cul1-F-box (SCF) family of E3 ubiquitin ligases that controls cell cycle through feedback mechanism affecting E2F1-3, it is acted upon itself by E2F1, 7/8, and the anaphase promoting complex (APC/C^Cdh1^) [33].

A slightly reduced frequency of the proliferation subset had direct co-expression association with CCNF in both COAD and LUAD (Tables 2 -3). The genes responsible for the reduction were ORC(s) 2/3/4/5, POLD4, and POLE4, suggesting origin licensing may not be properly regulated by CCNF1: Cdk1-3 in either tumor type.

### CCNJ, CCNO:Cdk2, CCNG1:Cdk4/5/6, CCNI:Cdk5

CCNJ knockdown in Drosophila leads to defects in chromosome segregation and progression through mitosis in early embryonic division cycles [34, 35]. Complete complexes necessary for origin licensing, firing, and DNA synthesis were not co-expressed fully with CCNJ, in agreement with a role for CCNJ downstream of most of these functions.

CCNO knockdown is associated with the lung disorder primary dyskinesia [36] [37]; it is expressed in multiciliate cells and it is thought to function in deuterosome mediated centriole amplification. CCNO:Cdk2 co-expression with the subset was partial in COAD, and deregulated with respect to proliferation in LUAD (Table 2-3).

CCNG1 overexpression in yeast leads to inhibition of pre-replicative complexes [38]; reflecting this finding CCNG1 has indirect co-expression with about two-thirds of the subset in LUAD (Tables 2-3). The lack of CCNG1 co-expression with proliferation in COAD has been previously observed [39].

CCNI is an atypical cyclin most abundant in postmitotic cells, it does not regulate proliferation [40]. Accordingly, CCNI had indirect co-expression with proliferation subset in COAD, and no co-expression correlation in LUAD; Cdk5 ranked directly co-expressed with a moderate frequency of proliferation factors (Tables 2-5).

### Transcriptional Cdks and their Cyclins

Canonical Cdks 1, 2, 4, and 6 are known as “Class I”, their function is to control cell cycle progression. Transcriptional Cdks (tCdks) 7, 8, 9, 12, 13, 19 make up “Class II”, they are thought to control pre-initiation complex assembly, initiation, elongation, and spliceosome recruitment in cyclical transcription oscillations [41]. Less well understood functionally, tCdks and their cyclin regulators form complexes CCNH: Cdk7, CCNC: Cdk 8/19 (Mediator complex components), CCNT1: Cdk 9, and CCNK: Cdk 9/12/13.

### CCNH: Cdk7, CCNK: Cdk9/12/13, CCNT1: Cdk 9

Cdk7 and Cdk9 are the most studied tCdks at present [41–43]; Cdk7 forms a regulatory complex with CCNH and MNAT1 that has CDK-activating kinase (CAK) activity on Cdk9 and other Cdks, and multiple roles in transcription. Cyclin regulator CCNH ranked direct in co-expression with many origin licensing factors in COAD, less in LUAD (Tables 2-3); the difference between the two tumor types was significant (p = 0.0106, Fisher’s exact test). Cdk7 ranked directly with far fewer of subset components in both COAD and LUAD (Tables 4-5). CCNT1: Cdk 9 complexes did not uniformly co-express highly with the subset (Tables 2-5). These data reflect a “non-cycling” expression pattern of the regulatory cyclins, and/or functional roles outside origin licensing, and firing.

CCNK: Cdk12/13 functions by phosphorylating RNA polymerase II C-terminal repeat domain. CCNK: Cdk12 is thought to regulate splicing and 3’ -end processing, knockdown in cell lines inhibits proliferation [44–47]. In accordance with knockdown experiments, CCNK directly co-expressed with some the proliferation subset in COAD, but not in LUAD suggesting deregulation (Tables 2-3); the difference between the two tumor types was significant (p = 0.0079, Fisher’s exact test). Cdk12 co-expressed directly with some of the subset in both tumors (Tables 4-5). Cdk13 unexpectedly indirectly co-expressed with the proliferation subset in COAD, less so in LUAD (Tables 4-5).

**Table 4.** Co-expression of Proliferation Genes with Cyclin Dependent Kinases in COAD.

4. Co-expression of Proliferation Genes with Cyclin Dependent Kinases in COAD
| Gene ID | Direct<br>co-expression<sup>1</sup><br>(%) | $r_s$<br>(range) | Indirect<br>co-expression<sup>1</sup><br>(%) | $r_s$<br>(range) | No<br>co-expression<br>(%) |
| --- | --- | --- | --- | --- | --- |
| CDK1 | 97 | 0.72 / 0.19 | 3 | -0.17 | 0 |
| CDK2 | 94 | 0.66 / 0.06 | 3 | -0.21 | 3 |
| CDK2AP1 | 6 | 0.21 / 0.17 | 3 | -0.19 | 91 |
| CDK2AP2 | 52 | 0.46 / 0.16 | 3 | -0.24 | 45 |
| CDK3 | 10 | 0.28 / 0.17 | 10 | -0.21 / -0.19 | 80 |
| CDK4 | 94 | 0.53 / 0.18 | 0 | 0 | 6 |
| CDK5 | 45 | 0.54 / 0.15 | 7 | -0.17 / -0.15 | 48 |
| CDK5R1 | 48 | 0.27 / 0.17 | 3 | -0.37 | 49 |
| CDK5R2 | 0 | 0 | 10 | -0.20 / -0.18 | 90 |
| CDK5RAP1 | 45 | 0.49 / 0.16 | 10 | -0.32 / -0.14 | 45 |
| CDK5RAP2 | 19 | 0.37 / 0.15 | 13 | -0.34 / -0.16 | 68 |
| CDK5RAP3 | 23 | 0.37 / 0.16 | 0 | 0 | 77 |
| CDK6 | 0 | 0 | 19 | -0.26 / 0.19 | 81 |
| CDK7 | 19 | 0.18 / 0.43 | 3 | -0.38 / -0.16 | 78 |
| CDK8 | 0 | 0 | 10 | -0.42 / -0.17 | 90 |
| CDK9 | 6 | 0.38 / 0.16 | 23 | -0.34 / -0.17 | 71 |
| CDK10 | 19 | 0.20 / 0.15 | 19 | -0.24 / -0.17 | 62 |
| CDK11A | 3 | 0.21 | 7 | -0.23 / -0.19 | 90 |
| CDK11B | 6 | 0.18 | 6 | -0.30 / -0.17 | 88 |
| CDK12 | 39 | 0.43 / 0.15 | 6 | -0.39 / -0.21 | 55 |
| CDK13 | 13 | 0.19 / 0.15 | 58 | -0.34 / -0.15 | 29 |
| CDK14 | 0 | 0 | 87 | -0.39 / -0.15 | 13 |
| CDK15 | 0 | 0 | 29 | -0.26 / -0.15 | 71 |
| CDK16 | 48 | 0.36 / 0.16 | 0 | 0 | 52 |
| CDK17 | 0 | 0 | 81 | -0.46 / -0.15 | 19 |
| CDK18 | 0 | 0 | 55 | -0.32 / -0.16 | 45 |
| CDK19 | 3 | 0.18 | 65 | -0.34 / -0.16 | 32 |
| CDK20 | 6 | 0.20 / 0.18 | 45 | -0.29 / -0.16 | 49 |
| CDKL1 | 32 | 0.31 / 0.16 | 7 | -0.17 | 61 |
| CDKL2 | 6 | 0.19 | 6 | -0.19 | 94 |
| CDKL3 | 0 | 0 | 52 | -0.32 / -0.18 | 48 |
| CDKL4 | 0 | 0 | 0 | 0 | 100 |
| CDKL5 | 3 | 0.18 | 71 | -0.44 / -0.14 | 26 |
| CDKAL1 | 23 | 0.44 / 0.15 | 3 | -0.28 | 74 |

<sup>1</sup>Percent of proliferation subset where Spearman Correlation ( $r_s$ ) ranks with significance, q value observed.
<sup>2</sup>Range ( $r_s$ ), most to least significant.

Comparing subcellular localization placed Cdk12 in the nucleus and Cdk13 in the nucleus, cytosol, extracellular space, and Golgi apparatus, suggesting it gets transported out of the nucleus and the plasma membrane; CCNK localizes predominantly to the nucleus [20]. Spearman correlations ranked “indirect” with proliferation factors and localization of Cdk13 outside of the nucleus suggest Cdk13 may not be involved, or is negatively involved, in proliferation in COAD and LUAD.

### CCNC: Cdk 8/19

CCNC and Cdk8 are components of the “Mediator complex” which bridges transcription factors at promoter DNA, at origin sites, enhancer or silencers, and RNA polymerase; it phosphorylates RNA Polymerase II and is required for the expression of all its transcripts [48]. Cdk8 is an oncogene in COAD, expression knockdown inhibits proliferation [11]. It is overexpressed in 42% of TCGA COAD and acts by increasing transcription of CTNNB1 and MYC, downstream effectors of *Wnt* signaling. By Spearman correlation, CCNC co-expressed highly with proliferation genes in COAD however Cdk8 did not, suggesting Cdk8 overexpression was not solely happening during S phase (Tables 2-5). An expression heatmap of the proliferation subset in comparison to CCNC and Cdk8 shows cases with Cdk8 over expression do not have uniformly high or low expression of proliferation factors (Figure 1A). Co-expression of CTNNB1 or MYC is significant and direct with Cdk8 in TCGA PANCAN COAD [11].

**Table 5.** Co-expression of Proliferation Genes with Cyclin Dependent Kinases in LUAD.

| Gene ID | Direct<br>co-expression<sup>1</sup><br>(%) | $r_s$<br>(range) | Indirect<br>co-expression<sup>1</sup><br>(%) | $r_s$<br>(range) | No<br>co-expression<br>(%) |
| --- | --- | --- | --- | --- | --- |
| CDK1 | 94 | 0.84 / 0.21 | 0 | 0 | 6 |
| CDK2 | 94 | 0.72 / 0.20 | 0 | 0 | 6 |
| CDK2AP1 | 58 | 0.40 / 0.16 | 3 | -0.23 | 39 |
| CDK2AP2 | 7 | 0.37 / 0.27 | 42 | -0.33 / -0.16 | 51 |
| CDK3 | 10 | 0.31 / 0.16 | 0 | 0 | 90 |
| CDK4 | 81 | 0.59 / 0.20 | 0 | 0 | 19 |
| CDK5 | 39 | 0.52 / 0.17 | 7 | -0.35 / -0.18 | 54 |
| CDK5R1 | 71 | 0.46 / 0.19 | 3 | -0.17 | 26 |
| CDK5R2 | 3 | 0.22 | 0 | 0 | 97 |
| CDK5RAP1 | 45 | 0.34 / 0.17 | 0 | 0 | 55 |
| CDK5RAP2 | 13 | 0.22 / 0.18 | 10 | -0.37 / -0.18 | 77 |
| CDK5RAP3 | 3 | 0.24 | 0 | 0 | 97 |
| CDK6 | 39 | 0.26 / 0.16 | 7 | -0.20 / -0.19 | 54 |
| CDK7 | 23 | 0.28 / 0.16 | 10 | -0.20 / -0.16 | 68 |
| CDK8 | 52 | 0.32 / 0.18 | 0 | 0 | 48 |
| CDK9 | 7 | 0.18 / 0.16 | 29 | -0.23 / -0.18 | 64 |
| CDK10 | 7 | 0.34 / 0.18 | 48 | -0.36 / -0.15 | 45 |
| CDK11A | 7 | 0.27 / 0.17 | 10 | -0.24 / -0.18 | 83 |
| CDK11B | 10 | 0.25 / 0.18 | 16 | -0.31 / -0.17 | 74 |
| CDK12 | 58 | 0.49 / 0.16 | 10 | -0.46 / -0.21 | 32 |
| CDK13 | 16 | 0.32 / 0.16 | 19 | -0.51 / -0.17 | 65 |
| CDK14 | 0 | 0 | 23 | -0.24 / -0.16 | 77 |
| CDK15 | 0 | 0 | 45 | -0.26 / -0.17 | 55 |
| CDK16 | 78 | 0.38 / 0.19 | 0 | 0 | 22 |
| CDK17 | 7 | 0.23 / 0.17 | 10 | -0.33 / -0.22 | 84 |
| CDK18 | 0 | 0 | 71 | -0.45 / -0.17 | 29 |
| CDK19 | 13 | 0.38 / 0.17 | 19 | -0.41 / -0.20 | 68 |
| CDK20 | 0 | 0 | 39 | -0.31 / -0.17 | 61 |
| CDKL1 | 0 | 0 | 65 | -0.35 / -0.18 | 35 |
| CDKL2 | 0 | 0 | 74 | -0.50 / -0.16 | 26 |
| CDKL3 | 6 | 0.29 / 0.17 | 6 | -0.22 / -0.20 | 88 |
| CDKL4 | 0 | 0 | 0 | 0 | 100 |
| CDKL5 | 3 | 0.27 | 78 | -0.50 / -0.15 | 19 |
| CDKAL1 | 23 | 0.39 / 0.20 | 3 | -0.27 | 74 |

<sup>1</sup>Percent of proliferation subset where Spearman Correlation ( $r_s$ ) ranks with significance, q value observed.
<sup>2</sup>Range ( $r_s$ ), most to least significant.

**Figure 1.**
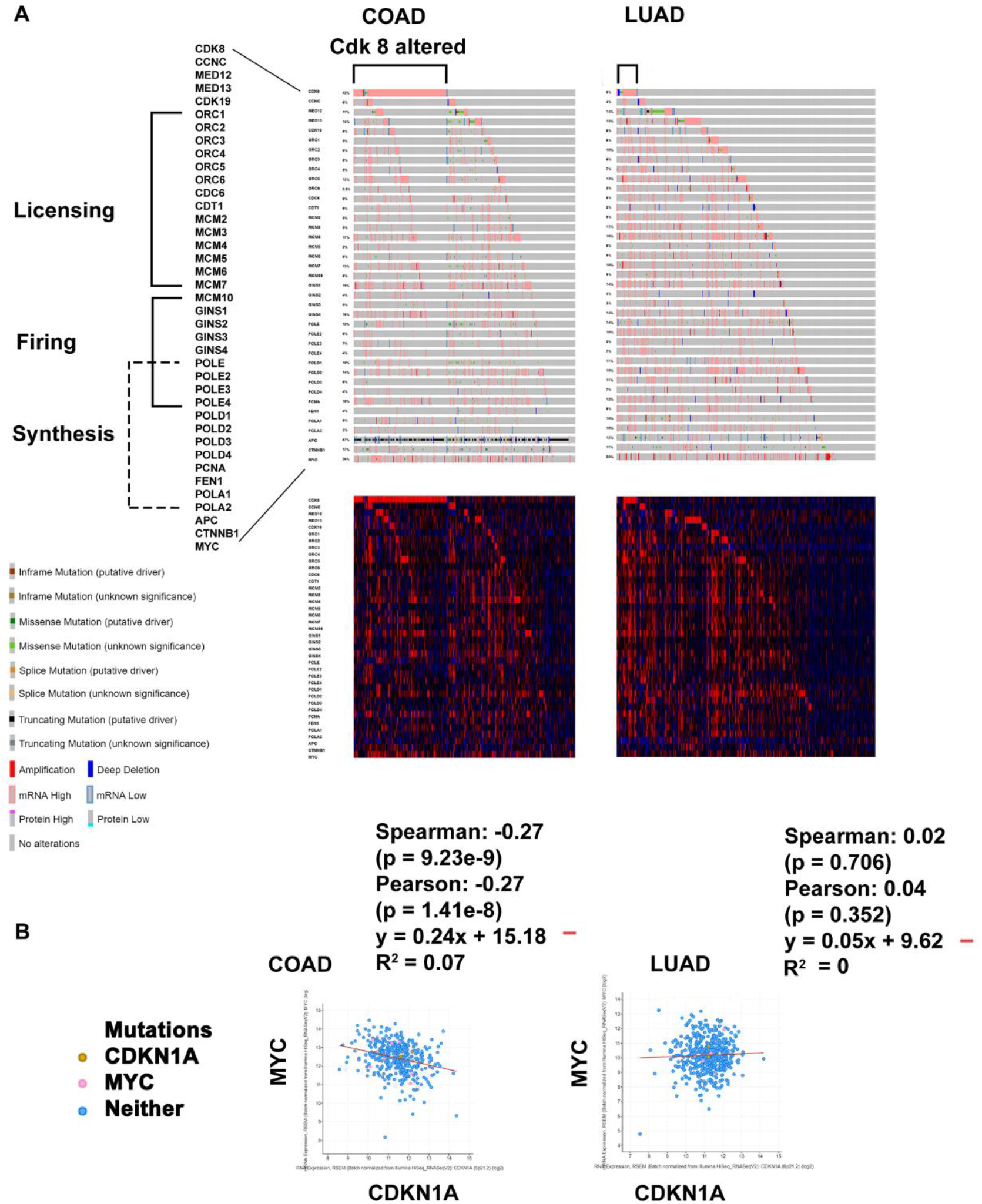
Genome and expression alterations to Mediator Complex components and proliferation subset factors. A, COAD (n = 439) and LUAD (n = 510) oncoprints from PanCan cohort. B, Co-expression of MYC versus CDKN1A in COAD and LUAD, with Spearman’s correlation.

CCNC and Cdk8 co-expressed directly with about half of the proliferation subset in LUAD (Tables 2-5); the difference in CCNC ranking between COAD and LUAD was significant (p = 0.0486, Fisher’s exact test). Overall alteration to the two genes in LUAD is approximately 4-8%.

### CCNY: Cdk14

Cdk 14 is highly expressed in brain [49], interacts with CCND3 and CDKN1A [50]; CDKN1A expression is controlled by TP53 (see below) [51]. Cdk14 is associated with *Wnt* signaling, peak levels of the membrane bound CCNY:Cdk14 complex occur during M phase, not S phase [12]. Accordingly CCNY had no significant co-expression with the proliferation subset in either COAD or LUAD, Cdk14 had high indirect co-expression with the subset in COAD (Tables 2-5). Overall alteration for both tumor types was 7-8% for CCNY, and 9% for Cdk14. CDKN1A Spearman correlation with MYC was examined in COAD and LUAD (Figure 1B), an indirect relationship was observed for COAD only.

Cdk14 over expression in relation to proliferation gene co-expression is shown by heatmap (Figure 2A). High Cdk14 correlates negatively with proliferation and positively with high hedgehog signaling (HH), and epithelial to mesenchyme transition (EMT) markers (Figure 2B); COAD cases with Cdk14 mutation in the protein kinase domain may reverse the correlation, proliferation co-expression high and HH and EMT low (Figure 2B) Other changes to *Wnt* signaling when Cdk14 was overexpressed were found for SFRP1/2/4 and DKK1-4 (Figure 2 C-D).

**Figure 2.**
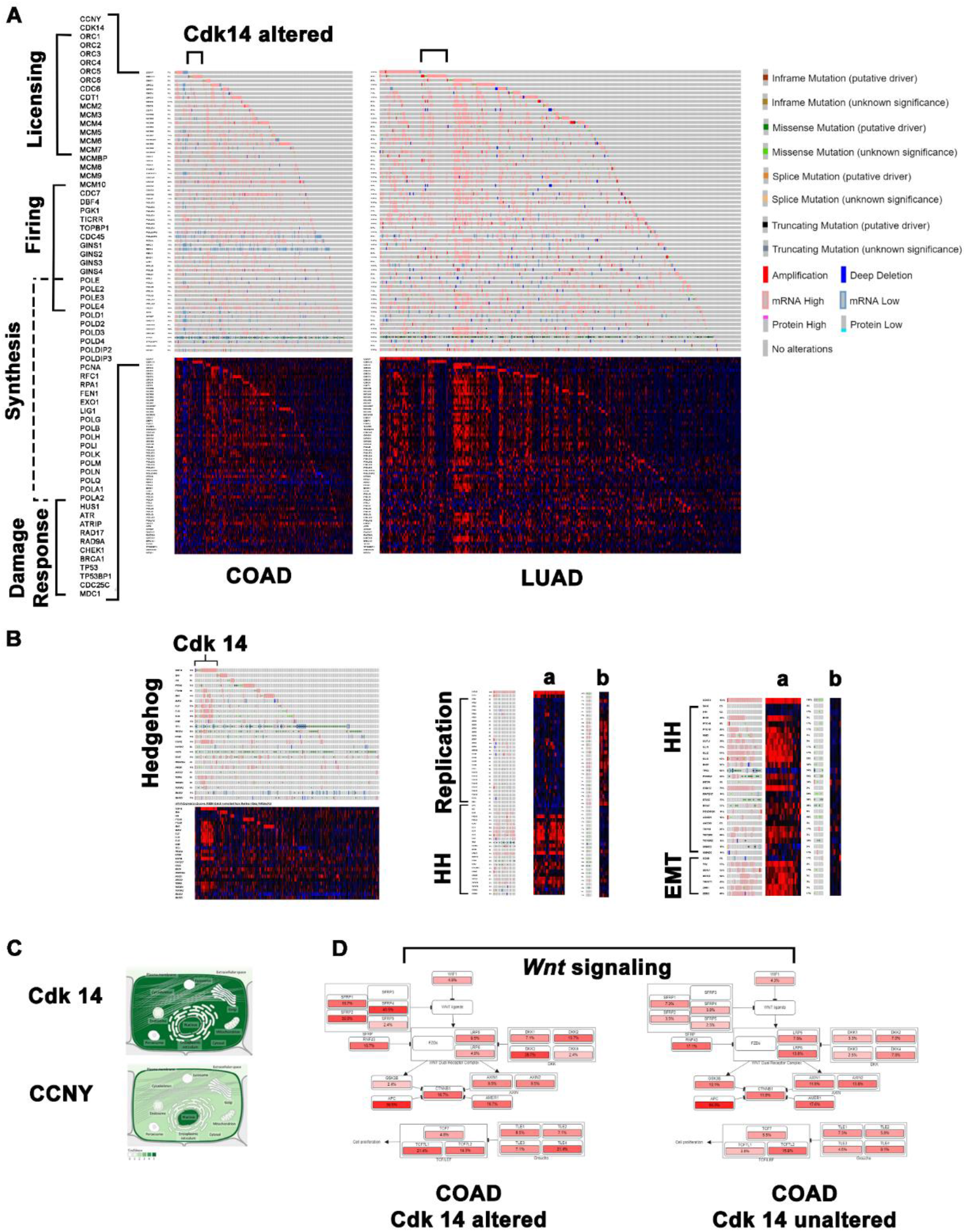
*Wnt* signaling components CCNY:Cdk14 relationship with proliferation. A, Oncoprint, PanCan cases with mRNA data (RNA Seq V2; COAD n = 592, LUAD n = 510). B, Oncoprint, Cdk 14 overexpressed COAD versus Hedgehog (HH) signaling pathway, replication, and epithelial to mesenchyme transition (EMT) markers. “a” Cdk 14 overexpressed, “b” Cdk 14 mutated. C, Subcellular localization [20]. *Wnt* signaling alterations, Cdk 14 overexpressed versus unaltered.

### Cdk Inhibitors

#### Cip/Kip Family, Universal Inhibitors

CDKN1A, (p21Cip/Waf1) is anti-proliferative, it binds with PCNA and suppresses G1/S and Rb phosphorylation. It is a component of Mediator complex, and functions in stem cells maintenance [52–55]. In this study some indirect correlation with the proliferation subset was seen in COAD (Table 7A), co-expression with MYC was indirect (Figure 1B) [14–16]. In LUAD [8, 9], significant indirect co-expression was not observed (Table 7B) nor was indirect co-expression with MYC (Figure 1B), suggesting deregulation.

**Table 6.**
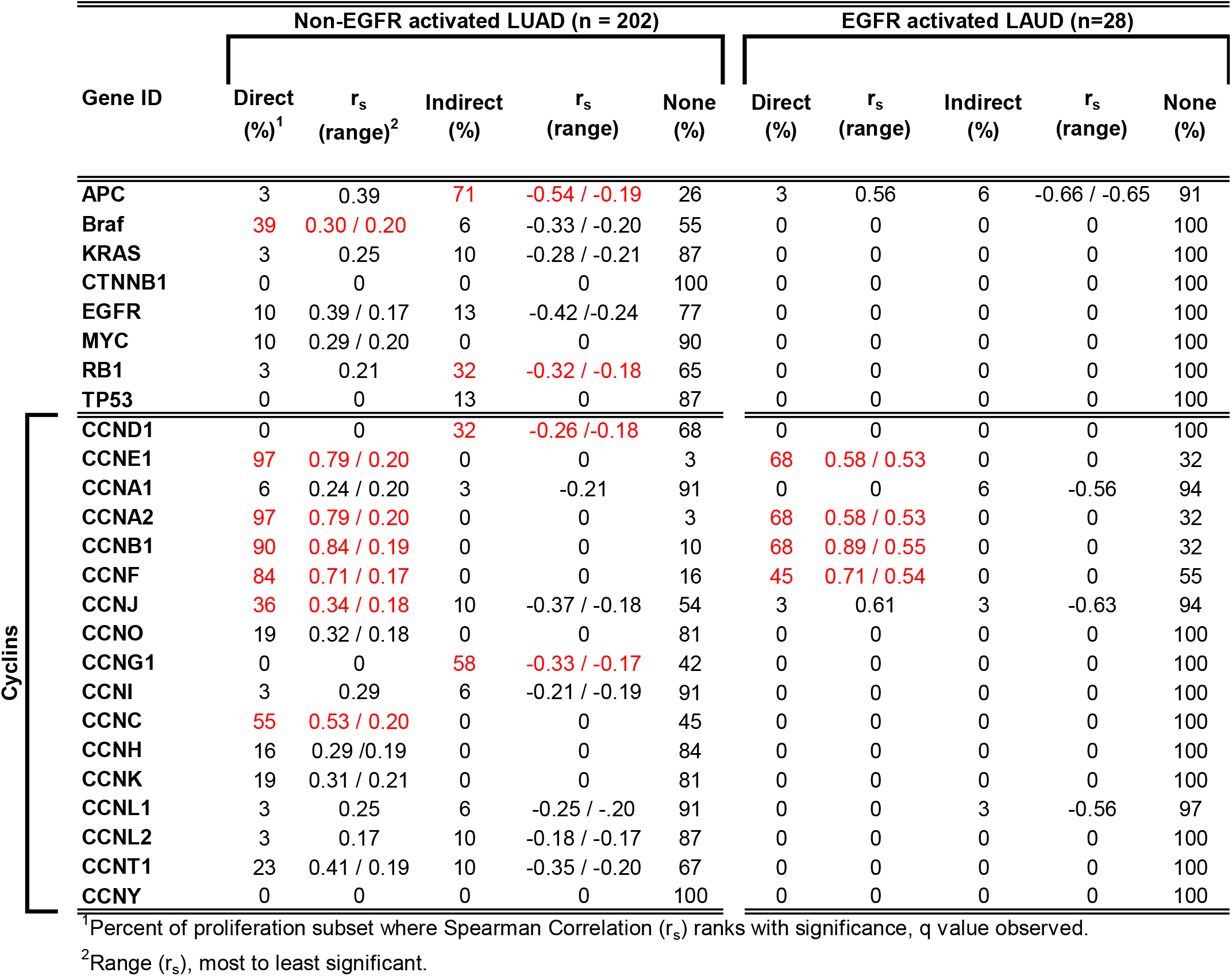
EGFR activated vs non-EGFR activated Lung Adenocarcinoma, Drivers and Cyclin co-expression with proliferation.

**Table 7.** Co-expression of Proliferation Genes with Cdk Inhibitors in COAD.

| Gene ID | Direct<br>co-expression<sup>1</sup><br>(%) | rs<br>(range) | Indirect<br>co-expression<sup>1</sup><br>(%) | rs<br>(range) | No<br>co-expression<br>(%) |
| --- | --- | --- | --- | --- | --- |
| CDKN1A | 3 | 0.25 | 35 | -0.37 / -0.15 | 62 |
| CDKN1B | 6 | 0.24 / 0.18 | 3 | -0.26 | 91 |
| CDKN1C | 0 | 0 | 74 | -0.30 / -0.16 | 26 |
| CDKN2A | 0 | 0 | 0 | 0 | 100 |
| CDKN2AIP | 6 | 0.23 / 0.18 | 0 | 0 | 94 |
| CDKN2B | 3 | 0.19 | 78 | -0.34 / -0.14 | 19 |
| CDKN2C | 10 | 0.35 / 0.24 | 13 | -0.33 / -0.28 | 77 |
| CDKN2D | 0 | 0 | 3 | -0.25 | 97 |
| CDKN3 | 94 | 0.81 / 0.14 | 0 | 0 | 6 |

**Co-expression of Proliferation Genes with Cdk Inhibitors in LUAD**
| Gene ID | Direct<br>co-expression<sup>1</sup><br>(%) | rs<br>(range) | Indirect<br>co-expression<sup>1</sup><br>(%) | rs<br>(range) | No<br>co-expression<br>(%) |
| --- | --- | --- | --- | --- | --- |
| CDKN1A | 0 | 0 | 7 | -0.21 / -0.18 | 93 |
| CDKN1B | 0 | 0 | 3 | -0.23 | 97 |
| CDKN1C | 0 | 0 | 26 | -0.34 | 74 |
| CDKN2A | 26 | 0.31 / 0.22 | 0 | 0 | 74 |
| CDKN2AIP | 0 | 0 | 55 | -0.30 / -0.19 | 45 |
| CDKN2B | 0 | 0 | 7 | -0.20 / 0.19 | 93 |
| CDKN2C | 68 | 0.35 / 0.16 | 3 | -0.26 | 29 |
| CDKN2D | 71 | 0.40 / 0.16 | 3 | -0.26 | 26 |
| CDKN3 | 97 | 0.82 / 0.18 | 0 | 0 | 3 |

<sup>1</sup>Percent of proliferation subset where Spearman Correlation ( $r_s$ ) ranks with significance, q value observed.
<sup>2</sup>Range ( $r_s$ ), most to least significant.

CDKN1B (p27KIP1) is anti-proliferative, MYC can inactivate the growth arrest it causes [56, 57]. In neither tumor type did CDKN1B co-express indirectly with the subset (Table 7A-B); nor did CDKN1B co-express indirectly with CCNE1. MYC significantly directly co-expressed with CDKN1B in COAD, not in LUAD [14–16].

CDKN1C (p57KIP2) shares homology with CDKN1A/B and is anti-proliferative causing G1 cell cycle arrest and binding PCNA in S phase, but it singularly functions in embryogenesis [58]. Diverse signals and transcription factors regulate its expression including micro-RNAs and lncRNAs, imprinting silences it in cancer; decreased expression is known to be inversely related to tumor growth in both tumor types. In this study, CDKN1C had a high inverse co-expression relationship to the subset in COAD, but not LUAD (Figure 7A-B). Total frequency alteration was 3-6% in COAD, 5-12% in LUAD.

### INK4 family, Inhibitors of Cdk4/6 and CCND1

CDKN2A isoforms, and CDKN2B inhibit Cdk4/6 and stabilize TP53 through MDM2. In COAD, CDKN2A did not co-express with the subset, CDKN2B indirectly co-expressed (Table 7A-B); genetic alteration for both genes was 5-7% of cases, mostly overexpression. In LUAD CDKN2A unexpectedly directly co-expressed with 26% of the subset. Interestingly 26% of LUAD had deep deletion or truncation mutations to the gene; mutations likely functionally permit the aberrant direction of coexpression [14–16]. CDKN2B had no co-expression with the subset, consistent with dysregulation; genetic alteration frequency to CDKN2B was 22%, mostly deep deletions.

Both CDKN2C and CDKN2D negatively regulate proliferation, CDKN2D has its highest transcriptional readout in S phase. In this study CDKN2C and CDKN2D co-expression with the proliferation subset was degregulated in COAD and LUAD (Table 7A-B).

### APC, CTNNB1, MYC, and KRAS

Apc is an antagonist of *Wnt* signaling [59–61]. MYC functions downstream of APC [2] and is involved in many biological processes and tumor types [62]. MYC and KRAS have been shown extensively to coordinate to influence proliferation and other processes [63].

In COAD, APC co-expression with the proliferation subset ranked “indirect” with high frequency; MYC and KRAS both ranked directly (Table 2). In LUAD, APC also co-expressed with the subset indirectly; MYC and KRAS direct co-expression was reduced (Table 3). Deregulation was more pronounced in EGFR activated LUAD over non EGFR activated LUAD (Table 6).

CTNNB1 binds APC anchoring the actin cytoskeleton to regulate contact inhibition, additionally acts as a transcriptional activator with TCF/LEF p300/CBP of *Wnt* signaling. CTNNB1 did not co-express with the subset in either COAD or LUAD, reflecting its known function (Tables 2-3).

To explore why APC would have the potential to act as a tumor suppressor in LUAD but not sustain mutation, transcriptional levels, mutations, and mutational signature of the gene were examined between the two tumor types. In COAD the most APC mutations are found in the last exon of the gene (exon 17) (Figure 3A-B), which codes for approximately 77% of the gene product. Transcriptional levels (RNA Seq V2 RSEM) of the APC gene were higher in COAD than in LUAD (p = 0.0182, unpaired t test/two tailed) (Figure 3C). The C to T mutational signature was significantly higher in COAD than in LUAD, (p = 0.0010, unpaired t test/two tailed) (Figure 3D). These results are addressed in the discussion section.

**Figure 3.**
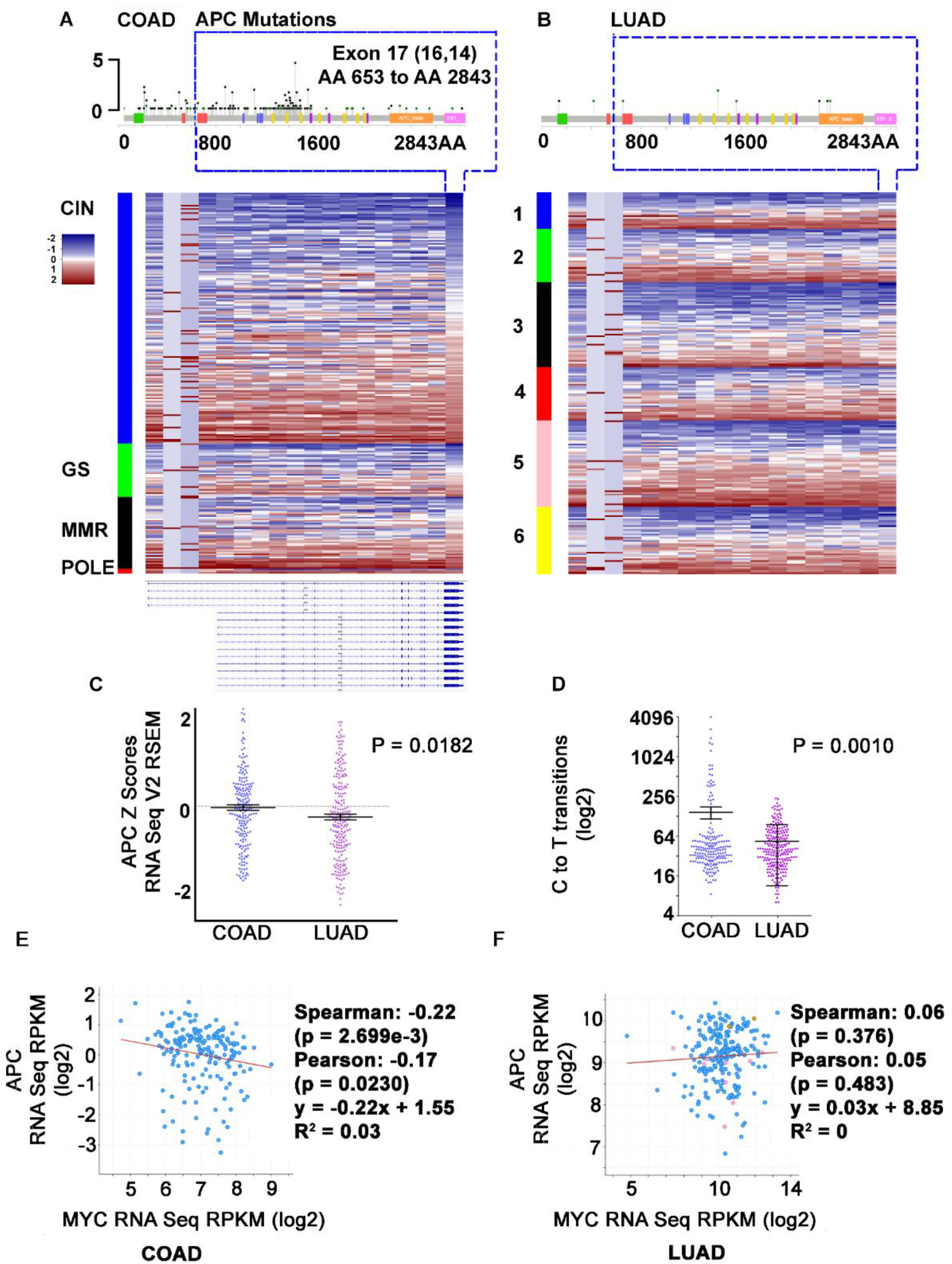
APC, COAD versus LUAD. A, APC genomic alterations in COAD, heatmap comparing APC mRNA expression for each exon (cases arranged in clusters: chromosome instabile “CIN”, genome stabile “GS”, Mismatch repair: “MMR”, POLE mutants), APC mRNA isoforms. B, APC mutations in LUAD, heatmap for each exon arranged in clusters 1-6. C, APC expression levels, COAD versus LUAD (significance by t test-two tailed). D, C to T transitions, COAD versus LUAD (t test-two tailed). E, Co-expression of MYC versus APC, COAD (complete samples) [1] and F, LUAD (complete samples) [8, 9].

*Wnt* signaling components SMAD(s) 2/3/4, and TCF7L2 also were found to have increased mutational frequency in COAD over LUAD, each with significant decreased co-expression with MYC (Figure 4A-B).

**Figure 4.**
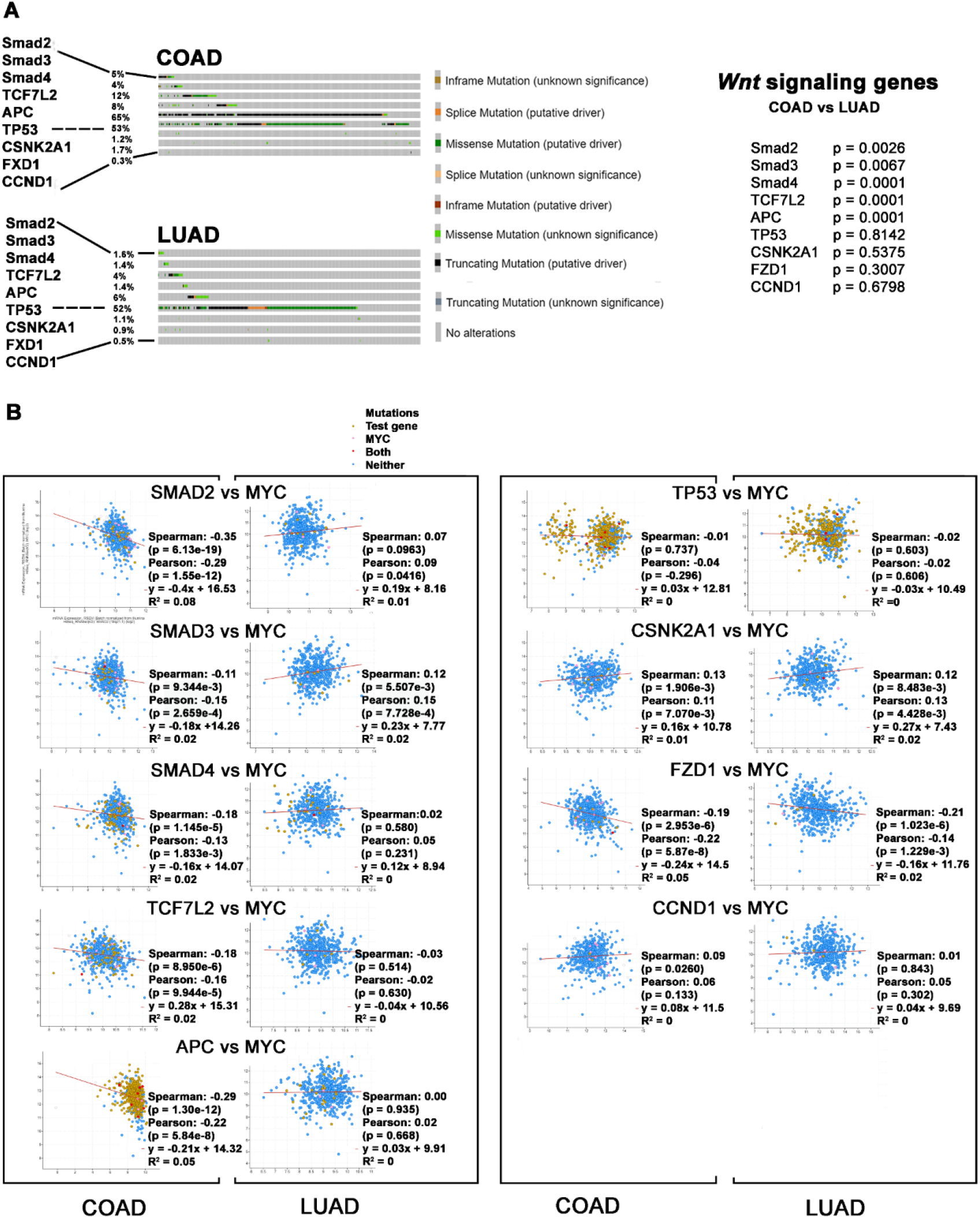
SMAD activators of *Wnt* signaling, mutations and expression relationship to MYC, COAD versus LUAD. A, Oncoprint (significance by Chi square analysis with Yates correction, see materials and methods).

### EGFR

LUAD separated into EGFR protein kinase activated versus EGFR nonactivated did not appreciably change the co-expression relationship of EGFR to the proliferation subset (Table 6). While EGFR is overexpressed in COAD, LUAD have most of the activation mutations (Figure 5A) many of which are due to exon 19 deletion; EGFR is more highly expressed in LUAD (Figure 5C) and mRNA isoforms exist (Figure 5D) (p = 0.0387, unpaired t test/two tailed). EGFR has a direct ranked co-expression relationship with MYC in COAD, and no co-expression relationship in LUAD (Figure 5E-F).

**Figure 5.**
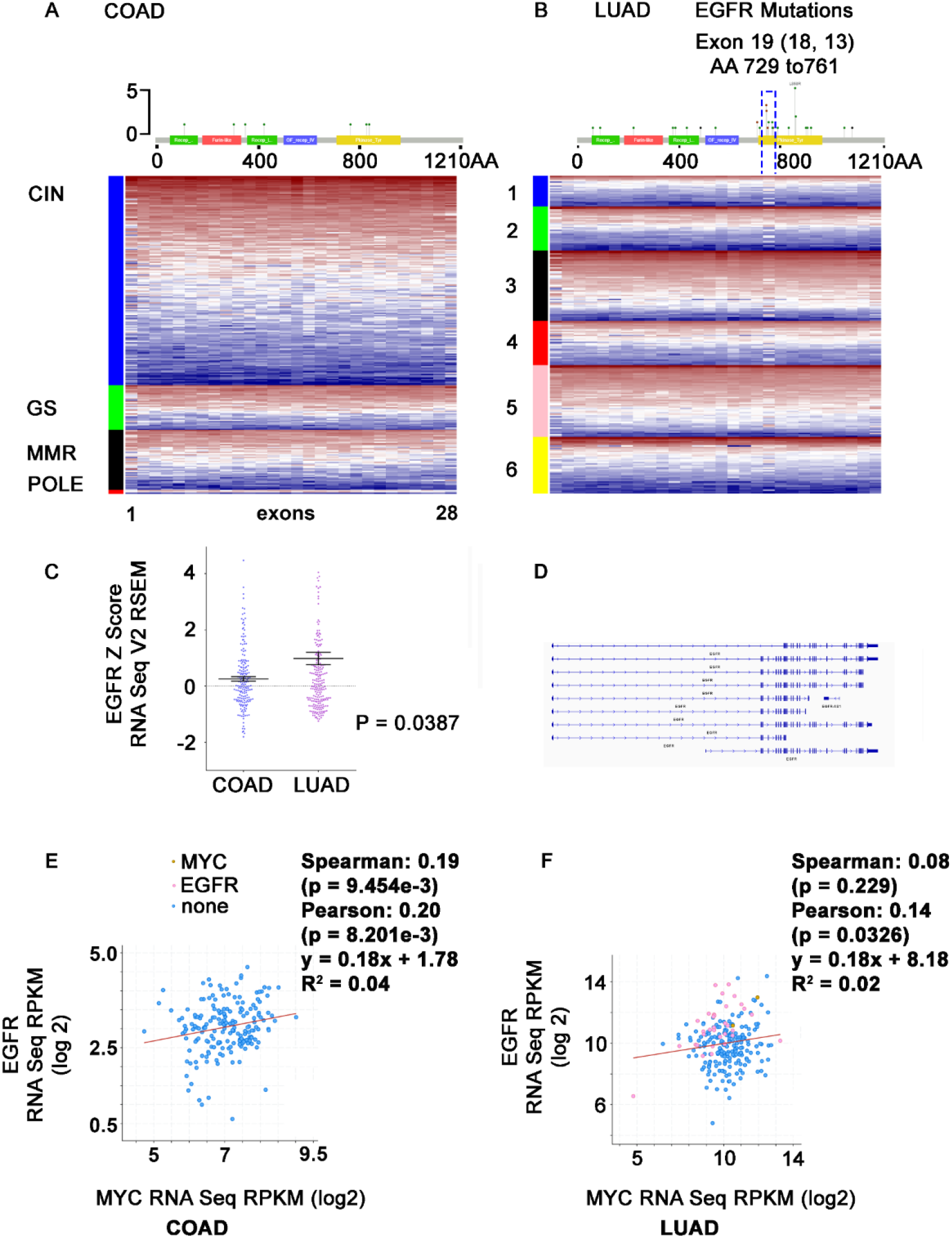
EGFR, COAD versus LUAD. A, genomic alteration in COAD, heatmap comparing mRNA expression for each exon. B, mutations in LUAD. C, EGFR mRNA expression levels (t test-two tailed). D, EGFR mRNA isoforms. E, Co-expression of MYC versus EGFR in COAD (complete samples) [1]. F, LUAD (complete samples [8, 9].

### E2F transcription factors and proliferation subset

E2F dysfunction in cancer has been recently reviewed [64]. E2F4/5/6 are canonical repressors that inhibit target gene expression by complexing with unphosphorylated pocket proteins (RB1, RBL1, and RBL2). Phosphorylation of pocket proteins leads to E2F4 and E2F5 (and E2F3B) removal from the complex, with E2F4/5 shuttled out of the nucleus releasing inhibition or target gene expression.

In this study E2F4/5/6 co-expression with proliferation factors is deregulated in EGFR activated tumors (Figure 7A), an earlier study showed EGFR activation correlates with decreased expression of proliferation factors [13].

### Braf Oncogene

Braf activates RAS/MAPK signaling to regulate growth and proliferation; in this study its expression neither “directly” nor “indirectly” co-expressed with the subset in

COAD or the total LUAD cohort (Tables 2-3). When EGFR activated cases were removed and assessed on their own, approximately 40% of the remaining subset (non-EGFR LUAD) had “direct” co-expression with Braf (Table 6).

## Discussion

Cell cycling depends on coordinated transcriptional oscillations, regulator/effector interactions and subsequent phosphorylation events. Genes responsible for these events are often pharmacological targets in cancer [65]. Spearman’s correlation compares the difference in expression ranking of two genes, and identifies the direction of expression relative to each other. The outcomes can be used to follow the transcriptional oscillations of the cell cycle, in this study they are used to associate genes with proliferation and S phase. The proliferation genes used as markers have known functions involved in origin licensing, firing, and integral DNA synthesis. DNA damage response and repair genes have been excluded, although some of the subset genes also function in these processes. Methylation and other epigenetic modifications which clearly play a role in coordinating transcription and proliferation and the definition of origin of replication sites, are also not examined in this study.

Canonical cyclins and the Cdks they regulate, by and large co-expressed with the proliferation subset as expected. During S phase for instance, when CCNE1 is highly expressed the highest frequency of proliferation factors in the subset are too as are Cdks 2 and 1. CCNA2:Cdk1/2 known to drive initiation and progression of proliferation and promote the beginning of mitosis [66] also highly co-expressed with the subset, as did maturation factor CCNB1. These observations are useful in verifying the system this study is based on since normal colon and lung expression data was not available through TCGA, they also support findings for genes with less established background.

Cell cycling is distinguished from the quiescence phase (G0) observed in most stem cells, by stages G1, S, G2, and M. G0 is defined as absence of proliferation with maintenance of proliferative potential [24]. Growth factors control progression from G0 to G1, S phase is marked by a restriction point after which growth factors have no effect. When examining CCNY:Cdk14, a membrane complex linked to *Wnt* signaling whose peak expression is thought during M phase [12, 67–69], Cdk14 overexpression without its CCNY regulator was found to correspond indirectly with proliferation gene expression and directly with co-expression of hedgehog signaling and epithelial to mesenchyme transition (EMT) genes, in COAD and most LUAD. The few COAD with Cdk14 mutations appeared to reverse the heatmap expression patterns for proliferation, hedgehog signaling, and EMT components. This behavior is reminiscent of the binary switch between G0 and proliferation; some evidence points to Cdk14 function in patterning and cell proliferation during embryonic development [70], regulator CCNY was identified by yeast two-hybrid screen [68]. Cigarette smoking may decrease Cdk14 levels in LUAD [71]. LUAD had more cases with high CCNY expression suggesting a mechanism for *Wnt* signaling activation in this proportion of the tumor type.

Progression from G1 into S phase occurs when retinoblastoma (Rb) is inactivated by phosphorylation, releasing E2F transcription factors. The relationship of E2Fs(1-8) with proliferation was examined to suggest Rb1 phosphorylation status *a priori*, as phosphorylation data for individual cases was not available. In LUAD, Rb1 alteration was mostly due to under expression, implying Rb1 was both inactivated and under expressed. E2Fs(1-8) co-expression ranked directly with a high frequency of proliferation factors, and mostly indirectly with Rb1 itself.

In EGFR activated LUAD, E2F(s)4/5/6 expression was completely deregulated with respect to S phase. E2Fs 4/5 normally function as repressors and are under Rb control, as it is inactivated by phosphorylation they get shuttled out of the nucleus as complexes are disrupted [64]. Knockdown experiments have suggested E2F6 controls progression through S phase, as S phase length is shortened [72]. However, E2F(s)4/5/6 were not mutated or aberrantly expressed (above or below the diploid tumor fraction), little evidence was found to confirm S phase timing alteration. Collectively, the data suggest EGFR protein kinase activation competes with Cyclin/Cdk complexes leading to inappropriate phosphorylation of RB1, RBL1,and RBL2 that prevents release of E2F(s)4/5 and subsequent cytoplasmic localization and proliferation. In a normal cell cyclin D:Cdk4/6, responsive to extracellular signals such as EGF, is thought to monophosphorylate Rb1 in early G1 at fourteen different sites which lead to preferential E2F binding patterns [25]. EGFR activated protein kinase probably interferes with this process, as well as having the potential to phosphorylate other substrates.

Another interesting consequence of removing EGFR activated cases from the LUAD cohort is the subsequent rise in the frequency of Braf direct ranked co-expression with genes of the proliferation subset. Mathematical removal of EGFR cases is a bit analogous to treating clinically with specific EGFR inhibitors, eliminating EGFR activated cells from tumor heterogeneity. The data suggest in many remaining cases Braf is more co-expressed with proliferation genes, at the same time KRAS and MYC become more deregulated with respect to S phase. Recent reports of a connection between Braf and EGFR acquired resistance mutations have been seen in several tumor types [73] [74]. Taken as a model, removal of cases with specific driver mutations and examination of the proliferation potential of remaining cases may be useful in identifying potential drivers of resistance where anti-tumor reagents exist; that EGFR and Braf are both members of the same signaling pathway (MAPK) may be significant. It may also suggest that synthetic lethality could be achieved by simultaneous use of inhibitors to both EGFR and Braf, if tolerable for the patient [75].

E2F(s)1-8 also co-expressed highly with proliferation factors in total COAD, however in this tumor type Rb1 was mostly over expressed. E2F(s) 3,5,6,7, and 8 all significantly directly co-expressed with Rb1 itself, implying release by Rb1 inactivation. Interestingly, E2F(s)1-8 in COAD had less direct ranked co-expression with proliferation then in LUAD (Figure 7A), possibly explaining why Rb knockout does not initiate intestinal cancer in mice [76]. Rb1, CCND1, and Cdk6 appeared deregulated with respect to proliferation in COAD.

### *Wnt* Signaling

*Wnt* based cell cycle regulation occurs through microtubule binding, positioning, and checkpoint mediated by APC, EB1, and GSK3B; signaling peaks during mitosis, as other processes stop [12]. APC is known to control proliferation through MYC [2], though the mechanism by which MYC carries out its role is not fully understood. In cell culture MYC over-expression promotes transformation, under-expression decreases proliferation and tumorigenesis. Complexing with MAX, MNT, CUBN, and MIZ1 which determine its function [77], it is transcribed during S phase and G2 [78]. Expression deregulation promotes mitotic problems causing apoptosis if the option is available, polyploidy if not. MYC protein domains imply it functions in the phosphodegron, chromatin remodeling, and transcription regulation. In animal models a MYC dominant negative mutation makes tumors driven by other genes (e.g. KRAS and TP53) regress, implying MYC regulates large numbers of genes that have never been fully identified.

Paradoxically most of the time experimental changes to MYC expression do not change mRNA levels of proposed downstream genes. Alternatively, MYC is now thought to control proliferation and tumorigenesis by functioning in basic transcription [78], that it frees promoters from stalled RNA polymerase complexes coordinating transcription elongation with cell cycle progression, preventing replication-transcription conflicts when “head to head” collisions occur between POLII and DNA polymerases, and restricting “R loop” formation.

In COAD APC co-expression with MYC was indirect, indicating maximum APC expression happens when MYC availability is at its lowest (Figure 3E). This occurs because MYC is transcribed during S phase, APC and other *Wnt* signaling components have high transcriptional readouts during M phase [12]. MYC was altered in twenty percent of the PANCAN LUAD cohort, over-expressed cases were completely deregulated with respect to proliferation subset co-expression [79] [14–16], and therefore S phase and the cell cycle.

This study suggests MYC is occasionally not available enough in COAD to carry out its elongation function in basic transcription which leads to mutation of the genes being transcribed. APC particularly lends itself to this model as most mutations (often towards the amino terminal of the protein) occur in exon 17, the last exon of the gene. *Wnt* signaling components SMAD(s) 2/3/4, and TCF7L2 also have significant increased mutational frequency in COAD over LUAD, each being expressed with decreased availability of MYC in COAD (Figure 4B). It is of interest that Braf driven LUAD are dependent on functional or activated *Wnt* signaling for initiation and maintenance [3–6], possibly achieved in a proportion by increased CCNY: Cdk14. The reduced mutational frequency of SMAD(s) 2/3/4, and TCF7L2 in the context of increased MYC in LUAD may not be happenstance. TP53, CSNK2A1, FZD1, CCND1 are all *Wnt* signaling genes whose expression is happening in both COAD and LUAD in a similar MYC (mostly direct) co-expression context. Unlike APC, SMAD(s) 2/3/4, and TCF7L2, the mutational signatures of these genes are not significantly different between the two tumor types. It may be relevant that COAD have high C to T transitions known to be a transcriptionally associated molecular signature [80], thought the result of faulty uracil incorporation into DNA. If MYC is not sufficiently around to carry out removal of R-loops for instance in COAD in *Wnt* signaling components, conjecturally one could have faulty uracil incorporation.

### MAPK Signaling

MYC is a target gene of both *Wnt* and MAPK signaling. MYC and KRAS involvement in cancer is thought to be co-operative and contextual genetically, how they co-operate is not fully understood. Their role in the immune response and STAT signaling is under investigation [63].

In this study KRAS and MYC examined together had high direct associated co-expression with proliferation genes in COAD, and low direct co-expression with the proliferation subset in LUAD; non-EGFR activated LUAD had further reduced direct co-expression. The similar common frequencies in each tumor type suggest underlying transcriptional coordination between MYC and KRAS, the difference in frequency between the tumor types implies a mechanism by which canonical MAPK signaling is deregulated.

KRAS and MYC are mutated roughly to the same extent in both tumor types and the changes are highly comparable (Figure 6A). A majority of KRAS mutations are to KRAS G12C/V/D, MYC is predominantly amplified with an occasional missense mutation. Although high expressors exist, MYC amplification in the genome does not generally lead to increased transcription in either tumor type. Some differences in alteration to other genes in the MAPK pathway exist (Figure 6B) between the two tumor types. For example, EGFR activation mutation often by exon 19 deletion are found in LUAD not COAD. Interestingly in COAD, EGFR has a direct co-expression relationship with MYC, and EGFR mutations are not found. More highly expressed in LUAD, the total EGFR cohort has no significant direct ranked co-expression with MYC (Figure 5F) and mutations are found. It may be of significance that EGFR activated cases locate predominantly above the EGFR/MYC Spearman correlation curve (Figure 5F, and Supplementary Figure 2A), suggesting somewhat higher EGFR transcription in those cases in lower MYC context (this is not identifiable examining median mRNA expression levels). EGFR does not significantly co-express directly with CCNK, which might imply splicing issues that could lead to the exon 19 deletion. This study suggests a major difference in MAPK signaling between COAD and LAUD is at the interface between KRAS and MYC transcription and their coordination with transcription of essential proliferation factors; perhaps implying involvement of tCdks and their regulators. Epigenetics probably has a role too in this process though, as mentioned earlier, it is not observed in this study.

**Figure 6.**
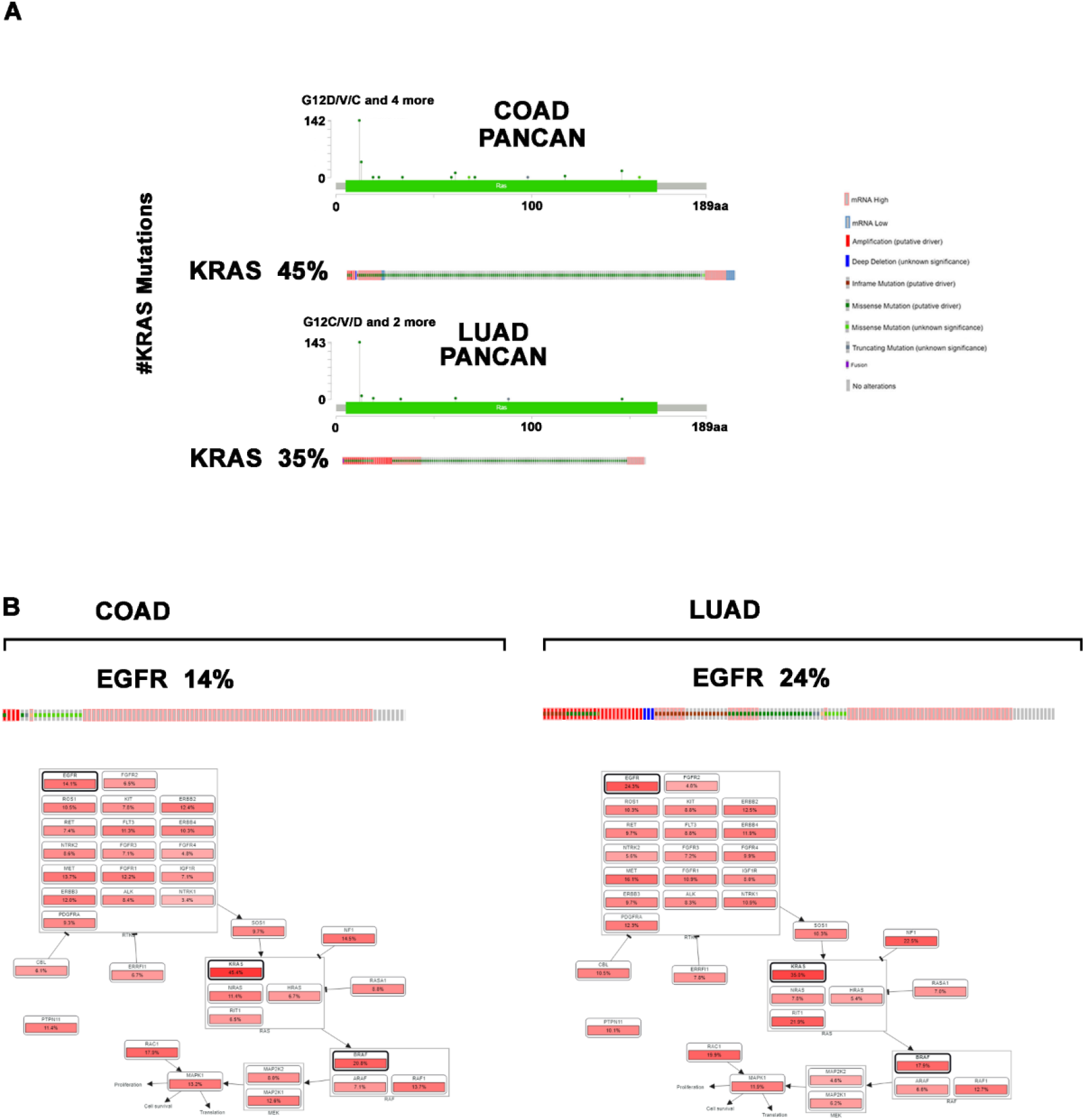
MAPK signlaing, COAD versus LUAD. A, KRAS mutations. B, EGFR mutations, and other MAPK components.

### Transcriptional Cyclins and tCDKs

CCNC, CCNH and CCNK are “non-cycling” cyclin regulators of tCdks, which generally explains their lack of a uniformly high direct rank co-expression with S phase proliferation factors. However, they do correlate with the decrease in direct rank co-expression with the proliferation subset between COAD and LUAD (Figure 7B). MYC, KRAS, CCNC, CCNH, and CCNK all have significant decreased frequency of direct ranked co-expression with essential proliferation factors in LUAD when compared to COAD. The tCdks the non-cycling cyclins regulate when examined alone have a more complicated relationship with the subset, but their activity is likely controlled less by their expression correlation than their cyclin regulator.

**Figure 7.**
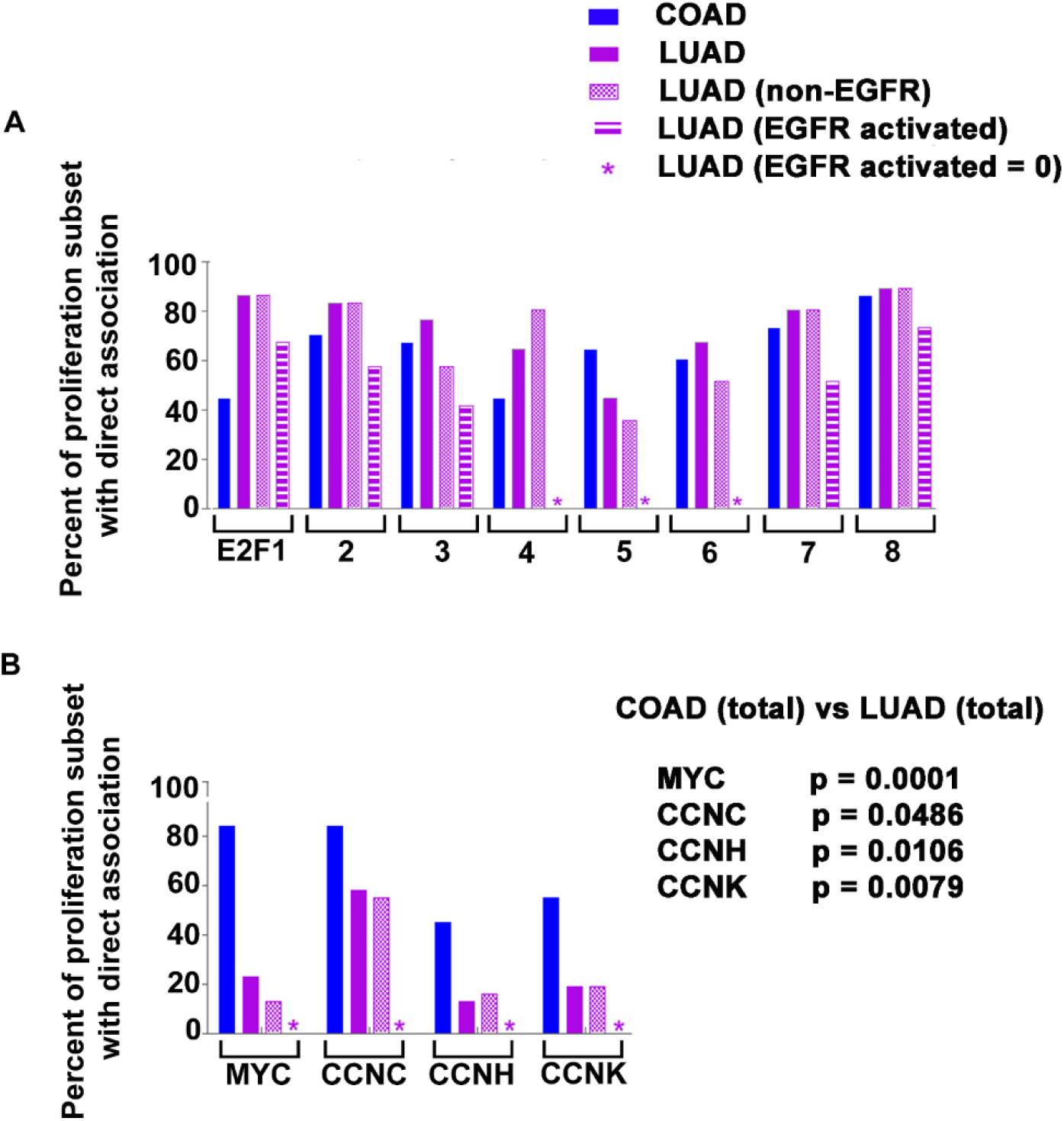
E2F, MYC, and regulator cyclin’s of tCdks relationship to proliferation. A, E2F 1-8 with direct co-expression to proliferation subset. B, MYC, CCNC, CCNH, and CCNK co-expression in COAD, LUAD, non-EGFR LUAD, and EGFR activated LUAD; significance by Fisher’s exact test.

## Conclusions

This study provides data that shows mechanisms responsible for the orchestration of expression between promoters enhancers and silencers of proliferation genes and the transcriptional transducers MYC/KRAS/BRAF/CCNC/CCNH/CCNK are different between COAD and LUAD. Since overall expression of these genes are comparable in both normal colon and lung, the disconnect between their co-expression with proliferation is almost certainly an integral part of the neoplastic process. The finding that MYC is indirectly co-expressed with APC, SMADS2/3/4, and TCF7L2 in COAD and not indirectly co-expressed in LUAD provides an explanation why those genes are susceptible to mutation in COAD and less so in LUAD. It may also be paradigmatic in explaining why mutations are difficult to find in some LUAD. MYC elevation of co-expression with respect to APC would be protective because enough MYC is present to carry out appropriate elongation function. Deregulation of MYC by elevation in LUAD contextually with other genes may lead to tumor progression directly, or alternatively be part of a shift to Braf coordination with proliferation leading to short circuiting of MYC and KRAS.

## Abbreviations

TCGA: The cancer genome atlas
PANCAN: Pan cancer
CDK: Cyclin dependent kinase
COAD: Colon adenocarcinoma
LUAD: Lung adenocarcinoma
*Wnt*: Wingless/integrated signaling
MAPK: Mitogen-activated protein kinase
APC: Adenomatous polyposis coli gene
CIN: Chromosomal instability
GS: Genome stabile
MMR: Mismatch repair
mRNA: Messenger RNA
RPKM: Reads per kilobase of transcription per million mapped reads
RSEM: RNA Seq by expectation maximization
FDR: False discovery rate
GTEx: Genotype-Tissue Expression portal
MPF: Maturation-promoting factor
SCF: Skp1-Cul1-F-box
HH: Hedgehog
EMT: Epithelial mesenchymal transition
RB: Retinoblastoma
EGFR: Epidermal growth factor receptor
r_s_ or ρ: Spearman’s correlation

## Declarations

### Ethics approval and consent to participate

The project protocol for the Harvard Genome Characterization Center (HGCC) was approved by the Brigham and Women’s Hospital Institutional Review Board. Access to TCGA controlled-access data was given by The Cancer Genome Atlas (TCGA) Data Access Committee. Participants agreed to participate and gave informed consent. Human Subjects Protection, Data Access Policies, and HIPAA Privacy Rule compliance were developed by the NCI and NHGRI to protect their privacy. This study is compliant with the TCGA “exclusivity period” for publication.

### Consent for publication

Not applicable.

### Availability of data and materials

The datasets generated and/or analyzed for the current study are available in cBioPortal [14–16] the Genome Data Commons[17] and Broad Institute [18].

### Authors’ contributions

MK was solely responsible for experimental conception and design, acquisition of data, and data analysis and interpretation, manuscript drafting of important intellectual content. The author gives approval of the final version for publication, takes responsibility for the content and agrees to be accountable for accuracy and integrity.

## Acknowledgements

The study was written during the pandemic of 2021, and the author wishes to express sincere appreciation to Dr. Kucherlapati for comments.

---

**Supplementary Figure 1.**
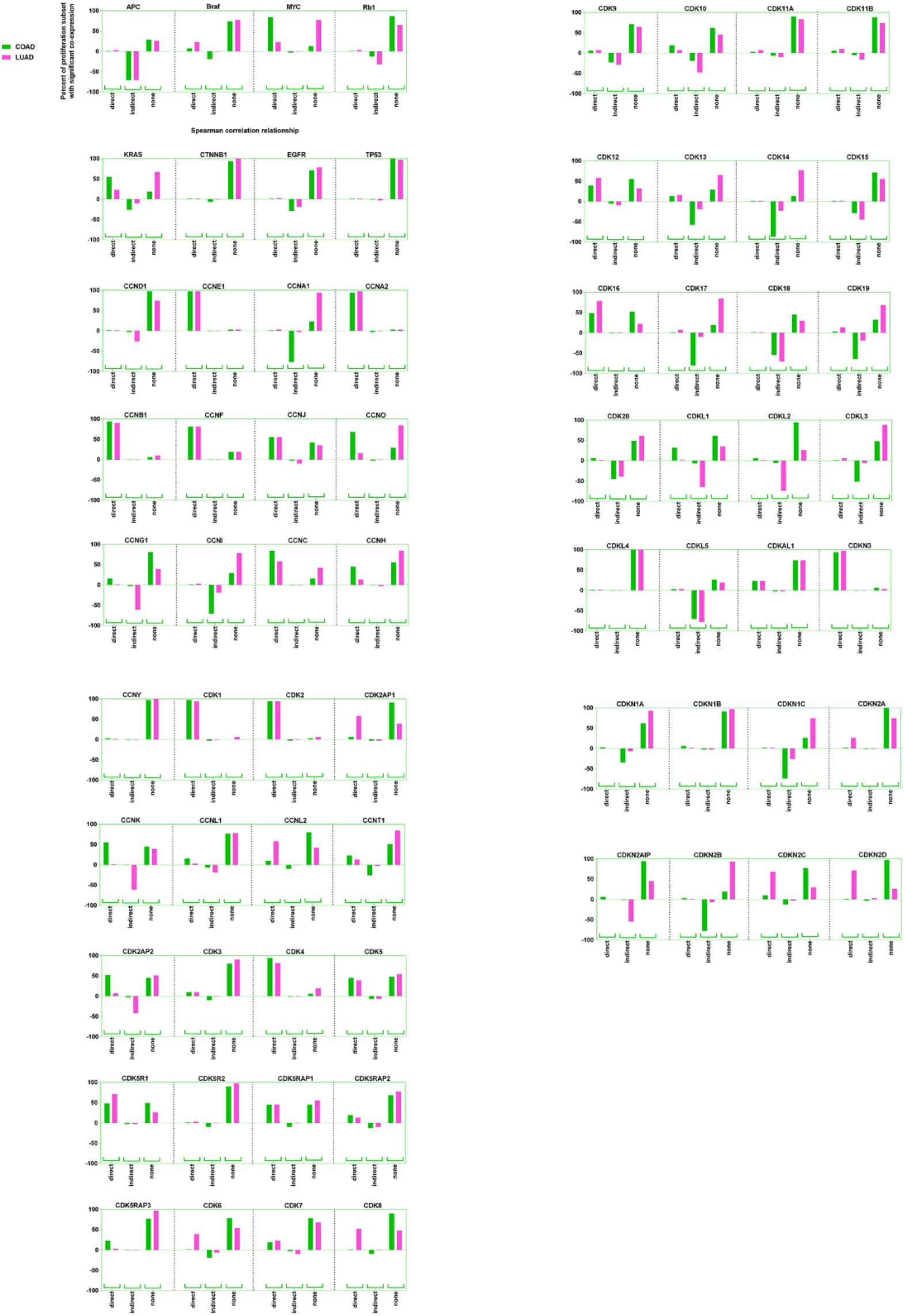
Data from Tables 2-7 given in graphical form.

**Supplementary Figure 2.**
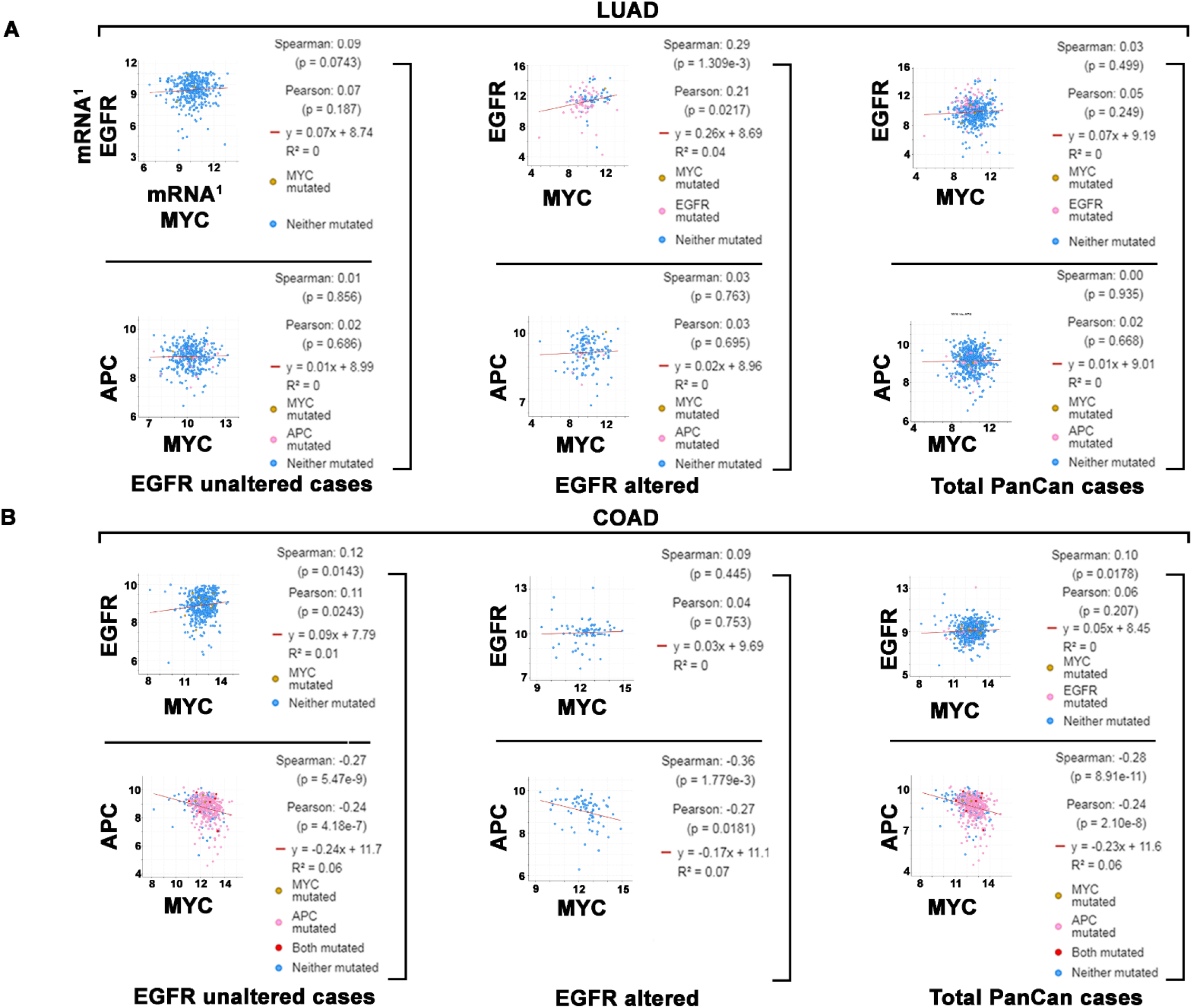
APC and MYC co-expression in EGFR cases from both COAD and LUAD PanCan data.

## REFERENCES

1. Cancer Genome Atlas N: Comprehensive molecular characterization of human colon and rectal cancer. Nature 2012, 487(7407):330–337.

2. He TC, Sparks AB, Rago C, Hermeking H, Zawel L, da Costa LT, Morin PJ, Vogelstein B, Kinzler KW: Identification of c-MYC as a target of the APC pathway. Science 1998, 281(5382):1509–1512.

3. Tammela T, Sanchez-Rivera FJ, Cetinbas NM, Wu K, Joshi NS, Helenius K, Park Y, Azimi R, Kerper NR, Wesselhoeft RA et al: A Wnt-producing niche drives proliferative potential and progression in lung adenocarcinoma. Nature 2017, 545(7654):355–359.

4. Juan J, Muraguchi T, Iezza G, Sears RC, McMahon M: Diminished WNT -> beta-catenin -> c-MYC signaling is a barrier for malignant progression of BRAFV600E-induced lung tumors. Genes & development 2014, 28(6):561–575.

5. Pacheco-Pinedo EC, Durham AC, Stewart KM, Goss AM, Lu MM, Demayo FJ, Morrisey EE: Wnt/beta-catenin signaling accelerates mouse lung tumorigenesis by imposing an embryonic distal progenitor phenotype on lung epithelium. The Journal of clinical investigation 2011, 121(5):1935–1945.

6. Stewart DJ: Wnt signaling pathway in non-small cell lung cancer. Journal of the National Cancer Institute 2014, 106(1):djt356.

7. Nguyen DX, Chiang AC, Zhang XH, Kim JY, Kris MG, Ladanyi M, Gerald WL, Massague J: WNT/TCF signaling through LEF1 and HOXB9 mediates lung adenocarcinoma metastasis. Cell 2009, 138(1):51–62.

8. Cancer Genome Atlas N, : Comprehensive molecular profiling of lung adenocarcinoma. Nature 2014, 511(7511):543–550.

9. Research TCGA: Author Correction: Comprehensive molecular profiling of lung adenocarcinoma. Nature 2018, 559(7715):E12.

10. Kim S, Xu X, Hecht A, Boyer TG: Mediator is a transducer of Wnt/beta-catenin signaling. The Journal of biological chemistry 2006, 281(20):14066–14075.

11. Firestein R, Bass AJ, Kim SY, Dunn IF, Silver SJ, Guney I, Freed E, Ligon AH, Vena N, Ogino S et al: CDK8 is a colorectal cancer oncogene that regulates beta-catenin activity. Nature 2008, 455(7212):547–551.

12. Kaldis P, Pagano M: Wnt signaling in mitosis. Developmental cell 2009, 17(6):749–750.

13. Kucherlapati MH: Modulation of proliferation factors in lung adenocarcinoma with an analysis of the transcriptional consequences of genomic EGFR activation. Oncotarget 2019, 10(65):6913–6933.

14. [http://www.cbioportal.org/index.do]

15. Cerami E, Gao J, Dogrusoz U, Gross BE, Sumer SO, Aksoy BA, Jacobsen A, Byrne CJ, Heuer ML, Larsson E et al: The cBio cancer genomics portal: an open platform for exploring multidimensional cancer genomics data. Cancer discovery 2012, 2(5):401–404.

16. Gao J, Aksoy BA, Dogrusoz U, Dresdner G, Gross B, Sumer SO, Sun Y, Jacobsen A, Sinha R, Larsson E et al: Integrative analysis of complex cancer genomics and clinical profiles using the cBioPortal. Science signaling 2013, 6(269):pl1.

17. [http://portal.gdc.cancer.gov/]

18. [http://gdac.broadinstitute.org/]

19. Wissler C: The Spearman Correlation Formula. Science 1905, 22(558):309-311.

20. [https://www.genecards.org/]

21. [https://www.graphpad.com/quickcalcs/contingency1/]

22. [www.graphpad.com]

23. Tchakarska G, Sola B: The double dealing of cyclin D1. Cell cycle 2020, 19(2):163–178.

24. Pennycook BR, Barr AR: Restriction point regulation at the crossroads between quiescence and cell proliferation. FEBS letters 2020.

25. Sanidas I, Morris R, Fella KA, Rumde PH, Boukhali M, Tai EC, Ting DT, Lawrence MS, Haas W, Dyson NJ: A Code of Mono-phosphorylation Modulates the Function of RB. Molecular cell 2019, 73(5):985–1000 e1006.

26. Bendris N, Loukil A, Cheung C, Arsic N, Rebouissou C, Hipskind R, Peter M, Lemmers B, Blanchard JM: Cyclin A2: a genuine cell cycle regulator? Biomolecular concepts 2012, 3(6):535–543.

27. Loukil A, Cheung CT, Bendris N, Lemmers B, Peter M, Blanchard JM: Cyclin A2: At the crossroads of cell cycle and cell invasion. World journal of biological chemistry 2015, 6(4):346–350.

28. Pagano M, Pepperkok R, Verde F, Ansorge W, Draetta G: Cyclin A is required at two points in the human cell cycle. The EMBO journal 1992, 11(3):961–971.

29. Wolgemuth DJ, Lele KM, Jobanputra V, Salazar G: The A-type cyclins and the meiotic cell cycle in mammalian male germ cells. International journal of andrology 2004, 27(4):192–199.

30. Liu D, Matzuk MM, Sung WK, Guo Q, Wang P, Wolgemuth DJ: Cyclin A1 is required for meiosis in the male mouse. Nature genetics 1998, 20(4):377–380.

31. Kitkumthorn N, Yanatatsanajit P, Kiatpongsan S, Phokaew C, Triratanachat S, Trivijitsilp P, Termrungruanglert W, Tresukosol D, Niruthisard S, Mutirangura A: Cyclin A1 promoter hypermethylation in human papillomavirus-associated cervical cancer. BMC cancer 2006, 6:55.

32. Porter LA, Donoghue DJ: Cyclin B1 and CDK1: nuclear localization and upstream regulators. Progress in cell cycle research 2003, 5:335–347.

33. Emanuele MJ, Enrico TP, Mouery RD, Wasserman D, Nachum S, Tzur A: Complex Cartography: Regulation of E2F Transcription Factors by Cyclin F and Ubiquitin. Trends in cell biology 2020, 30(8):640–652.

34. Kolonin MG, Finley RL, Jr.: A role for cyclin J in the rapid nuclear division cycles of early Drosophila embryogenesis. Developmental biology 2000, 227(2):661–672.

35. Atikukke G, Albosta P, Zhang H, Finley RL, Jr.: A role for Drosophila Cyclin J in oogenesis revealed by genetic interactions with the piRNA pathway. Mechanisms of development 2014, 133:64–76.

36. Terre B, Lewis M, Gil-Gomez G, Han Z, Lu H, Aguilera M, Prats N, Roy S, Zhao H, Stracker TH: Defects in efferent duct multiciliogenesis underlie male infertility in GEMC1-, MCIDAS- or CCNO-deficient mice. Development 2019, 146(8).

37. Ma JY, Ou-Yang YC, Luo YB, Wang ZB, Hou Y, Han ZM, Liu Z, Schatten H, Sun QY: Cyclin O regulates germinal vesicle breakdown in mouse oocytes. Biology of reproduction 2013, 88(5):110.

38. Tanaka S, Diffley JF: Deregulated G1-cyclin expression induces genomic instability by preventing efficient pre-RC formation. Genes & development 2002, 16(20):2639–2649.

39. Li JQ, Kubo A, Wu F, Usuki H, Fujita J, Bandoh S, Masaki T, Saoo K, Takeuchi H, Kobayashi S et al: Cyclin B1, unlike cyclin G1, increases significantly during colorectal carcinogenesis and during later metastasis to lymph nodes. International journal of oncology 2003, 22(5):1101–1110.

40. Brinkkoetter PT, Olivier P, Wu JS, Henderson S, Krofft RD, Pippin JW, Hockenbery D, Roberts JM, Shankland SJ: Cyclin I activates Cdk5 and regulates expression of Bcl-2 and Bcl-XL in postmitotic mouse cells. The Journal of clinical investigation 2009, 119(10):3089–3101.

41. Espinosa JM: Transcriptional CDKs in the spotlight. Transcription 2019, 10(2):45–46.

42. Fisher RP: Cdk7: a kinase at the core of transcription and in the crosshairs of cancer drug discovery. Transcription 2019, 10(2):47–56.

43. Bacon CW, D’Orso I: CDK9: a signaling hub for transcriptional control. Transcription 2019, 10(2):57–75.

44. Galbraith MD, Bender H, Espinosa JM: Therapeutic targeting of transcriptional cyclin-dependent kinases. Transcription 2019, 10(2):118–136.

45. Liang K, Gao X, Gilmore JM, Florens L, Washburn MP, Smith E, Shilatifard A: Characterization of human cyclin-dependent kinase 12 (CDK12) and CDK13 complexes in C-terminal domain phosphorylation, gene transcription, and RNA processing. Molecular and cellular biology 2015, 35(6):928–938.

46. Paculova H, Kohoutek J: The emerging roles of CDK12 in tumorigenesis. Cell division 2017, 12:7.

47. Greenleaf AL: Human CDK12 and CDK13, multi-tasking CTD kinases for the new millenium. Transcription 2019, 10(2):91–110.

48. Karijolich JJ, Hampsey M: The Mediator complex. Current biology : CB 2012, 22(24):R1030–1031.

49. Gao Y, Jiang M, Yang T, Ni J, Chen J: A Cdc2-related protein kinase hPFTAIRE1 from human brain interacting with 14-3-3 proteins. Cell research 2006, 16(6):539–547.

50. Shu F, Lv S, Qin Y, Ma X, Wang X, Peng X, Luo Y, Xu BE, Sun X, Wu J: Functional characterization of human PFTK1 as a cyclin-dependent kinase. Proceedings of the National Academy of Sciences of the United States of America 2007, 104(22):9248–9253.

51. el-Deiry WS, Tokino T, Velculescu VE, Levy DB, Parsons R, Trent JM, Lin D, Mercer WE, Kinzler KW, Vogelstein B: WAF1, a potential mediator of p53 tumor suppression. Cell 1993, 75(4):817–825.

52. Seleznik GM, Reding T, Peter L, Gupta A, Steiner SG, Sonda S, Verbeke CS, Dejardin E, Khatkov I, Segerer S et al: Development of autoimmune pancreatitis is independent of CDKN1A/p21-mediated pancreatic inflammation. Gut 2018, 67(9):1663–1673.

53. de Nooij JC, Hariharan IK: Uncoupling cell fate determination from patterned cell division in the Drosophila eye. Science 1995, 270(5238):983–985.

54. Mantel C, Luo Z, Canfield J, Braun S, Deng C, Broxmeyer HE: Involvement of p21cip-1 and p27kip-1 in the molecular mechanisms of steel factor-induced proliferative synergy in vitro and of p21cip-1 in the maintenance of stem/progenitor cells in vivo. Blood 1996, 88(10):3710–3719.

55. Xiao BD, Zhao YJ, Jia XY, Wu J, Wang YG, Huang F: Multifaceted p21 in carcinogenesis, stemness of tumor and tumor therapy. World J Stem Cells 2020, 12(6):481–487.

56. Amati B, Alevizopoulos K, Vlach J: Myc and the cell cycle. Frontiers in bioscience : a journal and virtual library 1998, 3:d250–268.

57. Vlach J, Hennecke S, Alevizopoulos K, Conti D, Amati B: Growth arrest by the cyclin-dependent kinase inhibitor p27Kip1 is abrogated by c-Myc. The EMBO journal 1996, 15(23):6595–6604.

58. Creff J, Besson A: Functional Versatility of the CDK Inhibitor p57(Kip2). Frontiers in cell and developmental biology 2020, 8:584590.

59. Oshima M, Oshima H, Kitagawa K, Kobayashi M, Itakura C, Taketo M: Loss of Apc heterozygosity and abnormal tissue building in nascent intestinal polyps in mice carrying a truncated Apc gene. Proc Natl Acad Sci U S A 1995, 92(10):4482–4486.

60. Shibata H, Toyama K, Shioya H, Ito M, Hirota M, Hasegawa S, Matsumoto H, Takano H, Akiyama T, Toyoshima K et al: Rapid colorectal adenoma formation initiated by conditional targeting of the Apc gene. Science 1997, 278(5335):120–123.

61. Sansom OJ, Reed KR, Hayes AJ, Ireland H, Brinkmann H, Newton IP, Batlle E, Simon-Assmann P, Clevers H, Nathke IS et al: Loss of Apc in vivo immediately perturbs Wnt signaling, differentiation, and migration. Genes Dev 2004, 18(12):1385–1390.

62. Li Y, Casey SC, Felsher DW: Inactivation of MYC reverses tumorigenesis. J Intern Med 2014, 276(1):52-60.

63. Mahauad-Fernandez WD, Felsher DW: The Myc and Ras Partnership in Cancer: Indistinguishable Alliance or Contextual Relationship? Cancer research 2020, 80(18):3799–3802.

64. Kent LN, Leone G: The broken cycle: E2F dysfunction in cancer. Nature reviews Cancer 2019, 19(6):326–338.

65. Suski JM, Braun M, Strmiska V, Sicinski P: Targeting cell-cycle machinery in cancer. Cancer cell 2021, 39(6):759–778.

66. Baumann K: Genome stability: Cyclin’ on mRNA. Nature reviews Molecular cell biology 2016, 17(11):676–677.

67. Davidson G, Shen J, Huang YL, Su Y, Karaulanov E, Bartscherer K, Hassler C, Stannek P, Boutros M, Niehrs C: Cell cycle control of wnt receptor activation. Developmental cell 2009, 17(6):788–799.

68. Jiang M, Gao Y, Yang T, Zhu X, Chen J: Cyclin Y, a novel membrane-associated cyclin, interacts with PFTK1. FEBS letters 2009, 583(13):2171–2178.

69. Ferguson FM, Doctor ZM, Ficarro SB, Browne CM, Marto JA, Johnson JL, Yaron TM, Cantley LC, Kim ND, Sim T et al: Discovery of Covalent CDK14 Inhibitors with Pan-TAIRE Family Specificity. Cell chemical biology 2019, 26(6):804–817 e812.

70. Zeng L, Cai C, Li S, Wang W, Li Y, Chen J, Zhu X, Zeng YA: Essential Roles of Cyclin Y-Like 1 and Cyclin Y in Dividing Wnt-Responsive Mammary Stem/Progenitor Cells. PLoS genetics 2016, 12(5):e1006055.

71. Pollack D, Xiao Y, Shrivasatava V, Levy A, Andrusier M, D’Armiento J, Holz MK, Vigodner M: CDK14 expression is down-regulated by cigarette smoke in vivo and in vitro. Toxicology letters 2015, 234(2):120–130.

72. Pennycook BR, Vesela E, Peripolli S, Singh T, Barr AR, Bertoli C, de Bruin RAM: E2F-dependent transcription determines replication capacity and S phase length. Nature communications 2020, 11(1):3503.

73. Prahallad A, Sun C, Huang S, Di Nicolantonio F, Salazar R, Zecchin D, Beijersbergen RL, Bardelli A, Bernards R: Unresponsiveness of colon cancer to BRAF(V600E) inhibition through feedback activation of EGFR. Nature 2012, 483(7387):100–103.

74. Stangl C, Post JB, van Roosmalen MJ, Hami N, Verlaan-Klink I, Vos HR, van Es RM, Koudijs MJ, Voest EE, Snippert HJG et al: Diverse BRAF Gene Fusions Confer Resistance to EGFR-Targeted Therapy via Differential Modulation of BRAF Activity. Molecular cancer research : MCR 2020, 18(4):537–548.

75. Xie Z, Gu Y, Xie X, Lin X, Ouyang M, Qin Y, Zhang J, Lizaso A, Chen S, Zhou C: Lung Adenocarcinoma Harboring Concomitant EGFR Mutations and BRAF V600E Responds to a Combination of Osimertinib and Vemurafenib to Overcome Osimertinib Resistance. Clinical lung cancer 2020.

76. Kucherlapati MH, Nguyen AA, Bronson RT, Kucherlapati RS: Inactivation of conditional Rb by Villin-Cre leads to aggressive tumors outside the gastrointestinal tract. Cancer research 2006, 66(7):3576–3583.

77. Carroll PA, Freie BW, Mathsyaraja H, Eisenman RN: The MYC transcription factor network: balancing metabolism, proliferation and oncogenesis. Frontiers of medicine 2018, 12(4):412–425.

78. Baluapuri A, Wolf E, Eilers M: Target gene-independent functions of MYC oncoproteins. Nat Rev Mol Cell Biol 2020, 21(5):255–267.

79. Little CD, Nau MM, Carney DN, Gazdar AF, Minna JD: Amplification and expression of the c-myc oncogene in human lung cancer cell lines. Nature 1983, 306(5939):194–196.

80. Owiti N, Stokdyk K, Kim N: The etiology of uracil residues in the Saccharomyces cerevisiae genomic DNA. Current genetics 2019, 65(2):393–399.

